# Exploring the role of the rich club in network control of neurocognitive states

**DOI:** 10.1101/2024.08.26.608239

**Authors:** Alina N. Podschun, Richard F. Betzel, Urs Braun, Sebastian Markett

**Affiliations:** Humboldt-Universität zu Berlin, Berlin, Germany; Indiana University Bloomington, Bloomington, IN, USA; Zentralinstitut für Seelische Gesundheit in Mannheim, Mannheim, Germany

## Abstract

We present a systematic analysis of the rich club as a possible control center of brain dynamics. Contrary to research stressing control properties of the rich club, our results indicate that the rich club and high-degree regions were not involved in optimally controlling the human connectome for a set of data-driven and task-evoked patterns of activity. Indeed, low-degree peripheral regions were more suited towards this. Findings were stable across group-level and individual-level rich club definitions, various cognitively meaningful brain state transitions, and different parameter settings. A region’s contribution to optimal control processes was associated with its affiliation with certain intrinsic connectivity networks and its position on the secondary, but not primary, cortical gradient. These results do not negate an integratory role of the rich club, but merely question its proposed role as a driver of control. Indeed, if it would inhabit such a role, we would have expected opposite results. Our findings fit with a position describing the rich club as a passive “data-highway” which, by means of its high connectivity, can be easily controlled by peripheral regions and thus facilitate relevant communication channels between them. We call for methodological expansions of the control theoretical toolbox allowing for elaborations on the temporal dynamics of control processes.

## Introduction

The human brain possesses the remarkable ability to adapt to changing demands by transitioning between various functional states (Shine et al., 2016; Wu et al., 2021). These brain states are carefully stabilized to support the current task or goal. When circumstances shift, the brain adapts by transitioning into more appropriate activity patterns (Benisty et al., 2024; Braun et al., 2021; Gonzalez-Castillo & Bandettini, 2018). Brain states are characterized by recurring activity patterns distributed across the brain, arising from both physiological and cognitive processes (Greene et al., 2023). Importantly, these patterns are believed to have direct influence on behavior (Benisty et al., 2024; Shine et al., 2016; Zink et al., 2021).

But where does this orchestration of neural activity originate? Emerging evidence suggests that the brain’s underlying structural network structure constrains dynamics of brain states (Betzel et al., 2016; Cole et al., 2016; Honey et al., 2007; Sorrentino et al., 2021; Suárez et al., 2020). Various brain regions communicate through a vast network of nerve fibers; adhering to certain organizational principles (Hagmann et al., 2008; Sporns & Betzel, 2016; Sporns & Zwi, 2004; Stiso & Bassett, 2018) that facilitate efficient, purposeful, and adaptive information processing (Rubinov, 2016; Rubinov & Sporns, 2010).

Furthermore, the brain’s governance is a collaborative effort rather than a traditional centralized control center (Betzel et al., 2016; Kim et al., 2018; Zink et al., 2021). One essential feature contributing to network-wide communication and integration is its rich club organization: This architecture feature involves well-connected hub regions that preferentially link to each other, creating a densely interconnected sub-network known as the “rich club” (van den Heuvel & Sporns, 2011). Although maintaining the rich club is costly due to the need for long-range and thick nerve fibers with high metabolic demand (Collin et al., 2014; van den Heuvel et al., 2012), it is most crucial for global integration and communication (de Reus & van den Heuvel, 2014; van den Heuvel et al., 2012).

Simulation studies have shown that such cortical hubs enable the brain to sustain a wide functional repertoire characterized by diverse dynamic configurations of peripheral regions (Bassett et al., 2013; Golos et al., 2015; Senden et al., 2014, 2017). These peripheral regions, with lower structural connectivity, interact with a stable high-degree core facilitated by the rich club, supporting rapid information exchange among relevant peripheral regions in a task-specific and goal-directed manner while remaining attentive to the entire network (Senden et al., 2018).

Network control theory, initially developed in engineering, offers a mathematical framework to quantify the energy required to maintain brain states and control transitions between them (Gu et al., 2015; Karrer et al., 2020; Parkes et al., 2024). The energy required for optimal transitions is related to the cognitive demand of specific states (Cornblath et al., 2020), with more demanding target states being more energetically costly (Braun et al., 2021; Luppi et al., 2023). Network control theory may thus be a suitable tool to quantify cognitive effort and executive function, assess impairments due to illness or injury, and to determine possible treatment routes (Dimulescu et al., 2021; Hahn et al., 2023; Jeganathan et al., 2018; Medaglia, 2019). Moreover, the application of network control theory identifies specific brain regions likely to control relevant transitions, often implicating regions high in network communicability (Gu et al., 2017), and high-degree regions within the rich club (Betzel et al., 2016; Braun et al., 2021). However, the specific role of the rich club in controlling transitions between behaviorally constrained brain states has yet to be explored.

In our current study, we leveraged network control theory to investigate regional optimal control contributions to brain state transitions across seven different tasks. We hypothesized that the rich club plays a pivotal role in network control, exerting a more substantial influence on brain state stabilization and transitions than expected by chance. Contrary to these expectations however, we found that rich club regions consistently contributed less to optimal control of brain state dynamics than a set of randomly chosen peripheral regions. This persisted across a diverse set of state transitions, connectome weighting schemes, and definitions of the rich club and reference sets. Instead, a region’s control impact was significantly related to membership in specific resting state networks, as well as its position on the secondary, but not primary, principal cortical gradient.

## Results

We studied the stability of brain states during seven canonical behavioral tasks from the Human Connectome Project (Van Essen et al., 2013, see Figure 1), and the control energy required to transition between these states. Using FA-weighted, undirected structural brain networks, we employed network control theory to approximate linear brain state dynamics, and the optimal control framework to determine the required control energy to maintain and switch between these task-constrained brain states.

**Figure 1.**
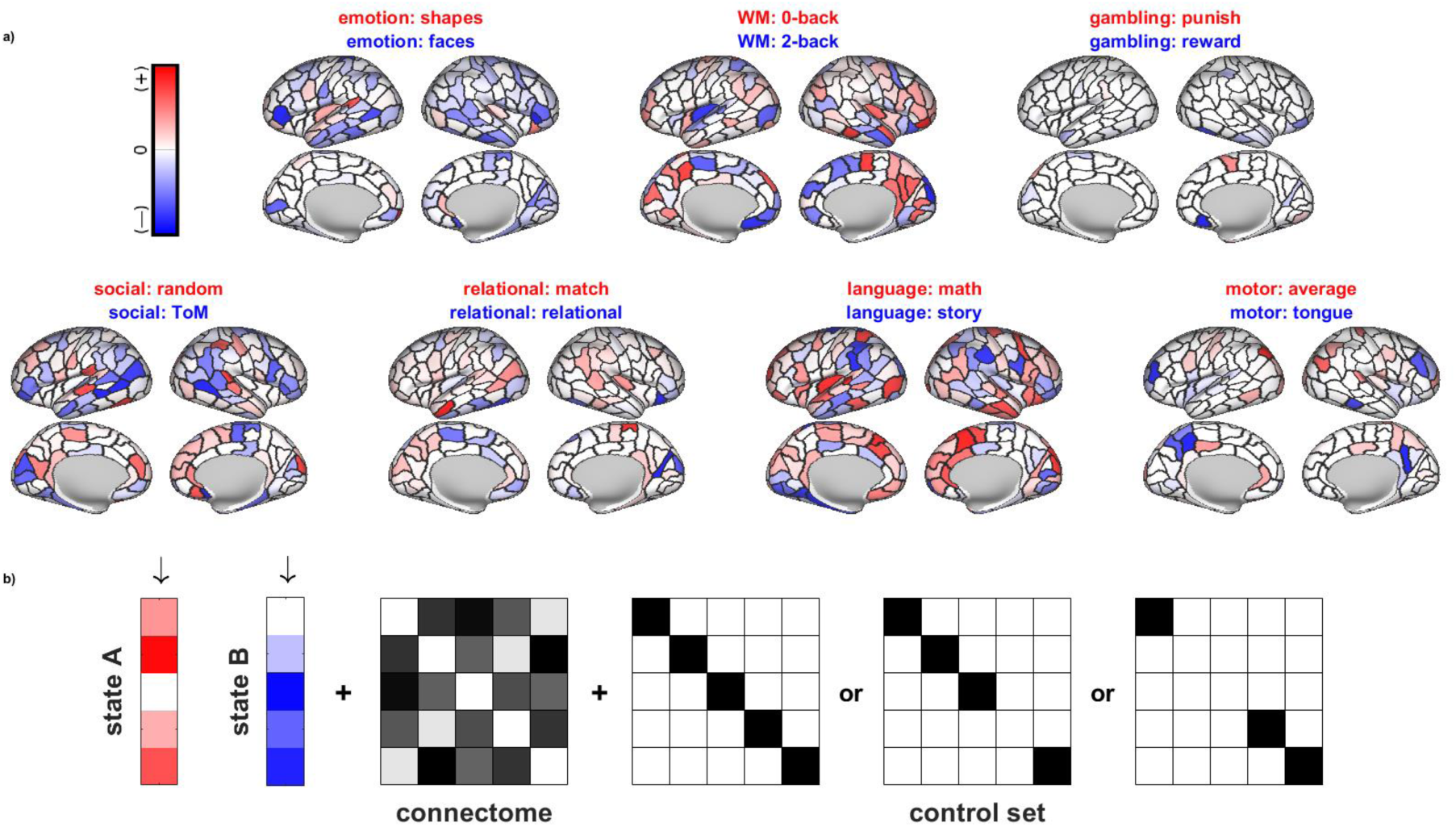
HCP brain states and overview over inputs into NCT analyses. We analyzed brain states constrained by the seven seven canonical HCP tasks. a) Each behavioral task consisted of a more stable (significant activity in red, usually the control condition) and a less stable brain state (significant activity in blue, usually the experimental condition). The more saturated the color, the higher the difference in activation between the two states. Figure 2 shows results of corresponding NCT analyses. b) Optimal control analyses require various inputs, here shown in a toy network consisting of five nodes. For each participant, trajectories between the two behaviorally constricted brain states were calculated constrained by a subject-specific, FA-weighted and undirected connectome and given a specific control matrix that specifies to which regions control is restricted. We investigated global control (all regions are allowed to exhibit control), regional control (all but one region are allowed to exhibit control) and also investigated the role of a subset of brain regions, excluding the rich club or a size-matched reference set from control. Given these components and penalizing as well as time parameters, control theoretical formulae can be solved for an optimized amount of control energy needed to traverse from the initial to the target state (see methods section).

### Global and regional control energy and stability

Each behavioral task is designed to elicit a minimum of two distinct brain states (see Figure 1). We first examined the global stability of all 14 brain states and the control energy needed to transition between the two states within each task, considering all brain regions as possible controllers. Except for the language task, the two brain states within each task differed significantly in their stability (all *p* < 0.01, Figure 2a). Transitions from the more stable to the less stable state required significantly more control energy than the reverse, again with the exception of the language task (all *p* < 0.05). Full ANOVA results can be found in the supplement (Supplemental Tables S1,S2).

**Figure 2.**
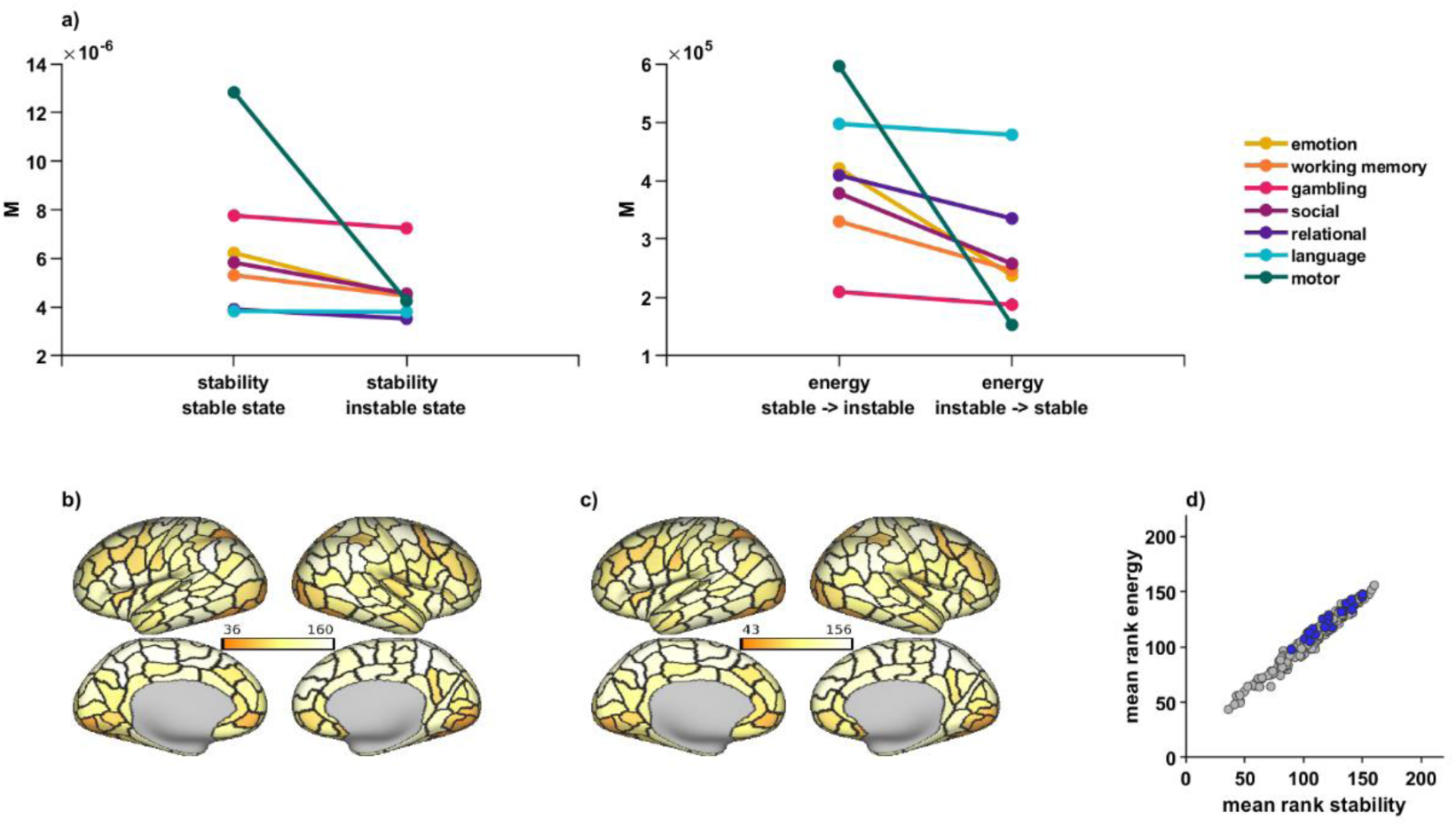
Results of global and regional control analyses - a subset of regions is task-generally implicated in control. We investigated global control of brain state trajectories, as well as possible task-general control roles of specific regions. a) For every task except for the language task, there was one state that was significantly more stable than the other state (left panel). Transitions from the stable state into the instable state were associated with significantly higher control energy needs than the reverse, again except for the language task (right panel). Displayed are mean global NCT values across individuals, and for each measure. b) - c) A subset of regions qualified as top controllers across tasks and was implicated both in maintenance of brain states (dark colors in b), as well as control of brain state transitions (dark colors in c). Another subset of nodes task-overarchingly did not contribute to regional control of brain states (light colors in b and c). Plotted is a region’s mean rank in regional control contribution across subjects and tasks, with dark colors indicating low ranks and thus high control contribution, light colors indicating high ranks and thus low control contribution. Black borders indicate different regions as defined in the Lausanne atlas. The supplement contains corresponding task- and measure-specific plots (Supplementary Figure 1). d) Shows the strong relationship between a region’s contribution to stability and energy measures. Rich club regions are indicated in blue; they overlapped with a set of comparatively high-ranking, i.e. not impactful, regions.

In a second step, we quantified each individual brain region’s contribution to maintaining and transitioning between brain states by estimating the change in global control stability and energy when excluding each region from the control matrix (following Braun et al., 2021). In order to determine whether individual brain regions function as task-general control nodes, we rank-ordered regions by their control impact for each task; and averaged each region’s rank for both stability and energy measures across all tasks.

Brain regions contributing to task-general brain state maintenance also facilitated task-general brain state transitions (Figure 2b-d). For stability measures, we observed mean ranks between rank_min_ = 36 and rank_max_ = 160 across tasks. For transition measures, we observed mean ranks between rank_min_ = 43 and rank_max_ = 156 across tasks. With lower ranks indicating higher control impact, minima values show that while some regions had more task-general contributions than others, there was still considerable variation across tasks and measures in top control nodes. Maxima values, however, point to a subset of nodes that did consistently not contribute to the control of brain states across tasks.

### Rich club regions have a smaller impact on control metrics than expected by chance

We identified 21 group-level rich club regions, which were most consistently assigned to the rich club based on the maximal normalized rich club coefficient at the subject-level (Riedel et al., 2022). These regions included the superior parietal cortex, insula, precuneus, rostral middle frontal cortex, superior frontal cortex bilaterally as well as *left* pericalcarine cortex, lateral occipital cortex, pars opercularis, inferior parietal cortex and *right* pars orbitalis and temporal pole (Figure 3a,c).

**Figure 3.**
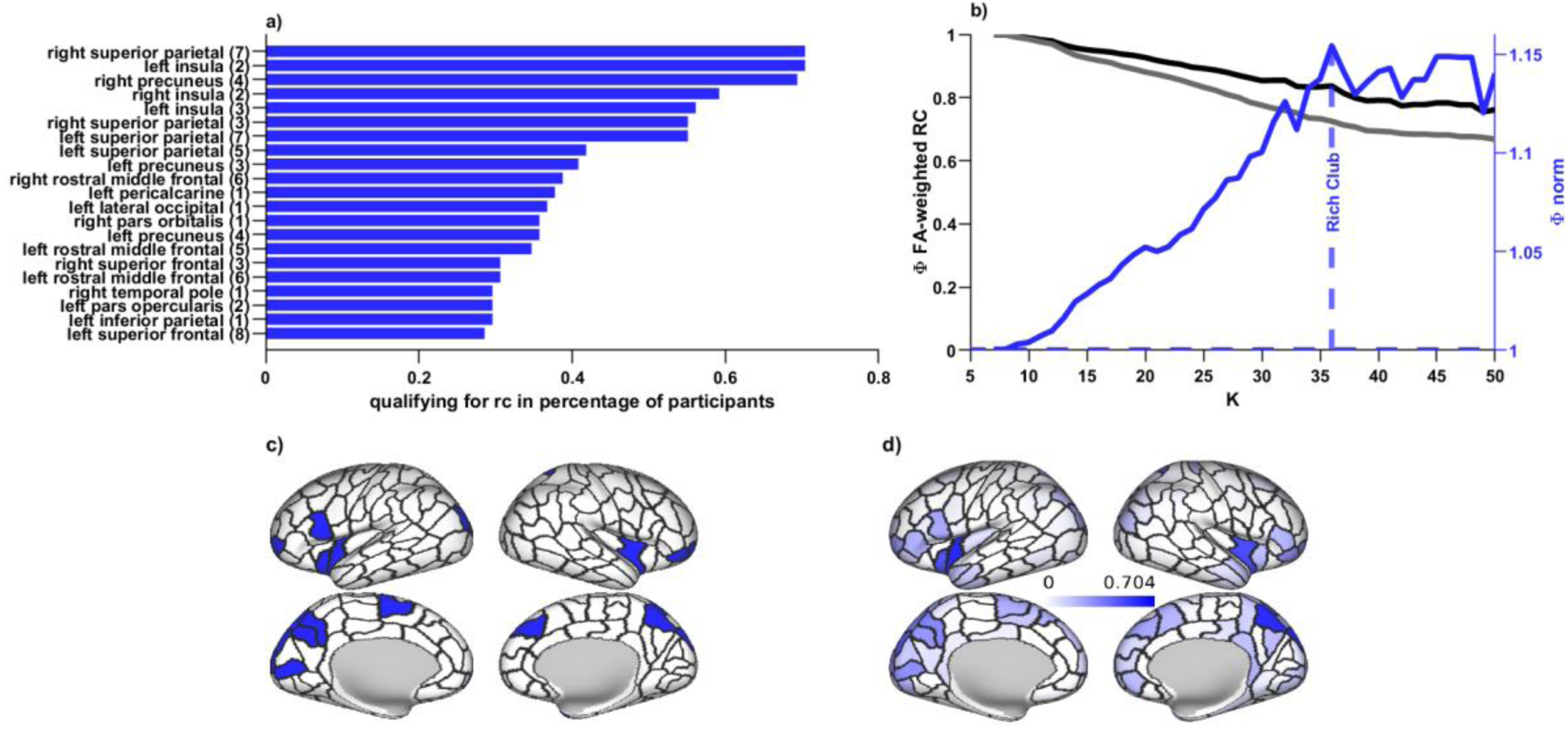
Rich-clubness and the group-level rich club. Visualizations of rich-clubness and our group-level rich club. a) 21 regions were qualifying for the group-level rich club and are shown here; corresponding relative frequency with which a given region was assigned rich club membership on a subject-specific level is indicated in the bars and varied greatly. b) The rich club was defined by comparing the structural connectivity of the empirical network (black) in relation to a randomized network (gray) for every nodal degree level k. Comparison of the two leads to a normalized (blue) rich club curve, which peaks at *k_max(Φ)_*. This peak characterizes the degree level k at which connectivity in the empirical network is maximally stronger than expected by chance, and was set as the starting point of the rich club, with all regions with nodal degree greater than *k_max(Φ)_* being considered rich club members. Curves are based on a group-level connectome and included merely for visualization. c) This procedure lead to 21 group-level rich club members (see also a), which are here projected onto an inflated map of the cortical surface. d) There was considerable interindividual variation with which a region would be assigned rich club status on a subject-specific level. Saturation of the color indicates the relative frequency with which a region would be assigned rich club membership on an individual level; the 21 regions for which this was most frequently the case constitute the group level rich club shown in a) and c).

Rich club regions spatially overlapped with middle to high rank on the task-general control metric, indicating negligible control impact across tasks (Figure 2d, compare also Figure 3c to Figures 2b,c). We ascertained this observation through a spatial permutation test using the spin-test procedure, which shuffled brain regions across the cortex while preserving the cortex’s inherent autocorrelation structure. Rich club regions had higher ranks for both stability (*M_rank_* = 122; *p* = 0.0329) and energy (*M_rank_* = 123; *p* = 0.0069) than expected by chance (10,000 permutations), indicating that rich club regions indeed contributed less to stability and control energy than the periphery.

In order to ensure that this observation was not confounded by the strict binary division of brain regions into a rich cub and a periphery (Alstott et al., 2014; Griffa & Van den Heuvel, 2018), we additionally correlated regional nodal degree with the regional ranks, again comparing the resulting correlation to a null distribution of 10,000 permutations using a spin test. Nodal degree correlated positively with a region’s rank for both stability (*r* = 0.35; *p* < 0.0001) and control energy (*r* = 0.42; *p* < 0.0001) measures, confirming high degree regions contributed less to both state maintenance as well as transitions between states.

While our group-level rich club included those 21 regions most frequently considered rich club members in subject-level analyses, there was still considerable variation in the relative frequency by which a region would be assigned to the rich club (Figure 3d). To ascertain that our observation of negligible control contribution by the rich club was not a confound of the group-level rich club definition, we repeated all analyses based on individual-level rich club assignments, where we defined the 21 top-degree regions in each participant as their individual rich club. These analyses confirmed all group-level results: None of the participants exhibited a significant involvement of rich club regions in stabilizing or transitioning between brain states. In the majority of participants, rich club regions contributed significantly less to both brain state stability (73.47%) and transitions (85.71%) than expected by chance. For a subset of subjects with comparatively low-ranking (i.e.: comparatively impactful) rich clubs, findings were non-significant; for this subset of participants, rich club nodes and peripheral nodes did not differ in their control contribution (see Supplementary Figure 2).

### Excluding the rich club from the control matrix leads to a significantly smaller impact on control metrics than excluding a size-matched set of random regions

We further corroborated our finding through optimal control analyses in which we excluded all rich club regions from the matrix of possible control regions. This exclusion resulted in lower control energy requirements for stabilizing (all *p* < 0.01, Figure 4) and transitioning between brain states (all *p* < 0.05, Figure 4a) in all seven tasks, compared to excluding a random group of 21 regions (spin test with 100 permutations). These findings again confirm that - compared to any other group of brain regions - the rich club contributed less to the stabilization of brain states and less to the controlled transitions between states (see Supplementary Tables S3 and S4 for detailed statistics).

**Figure 4.**
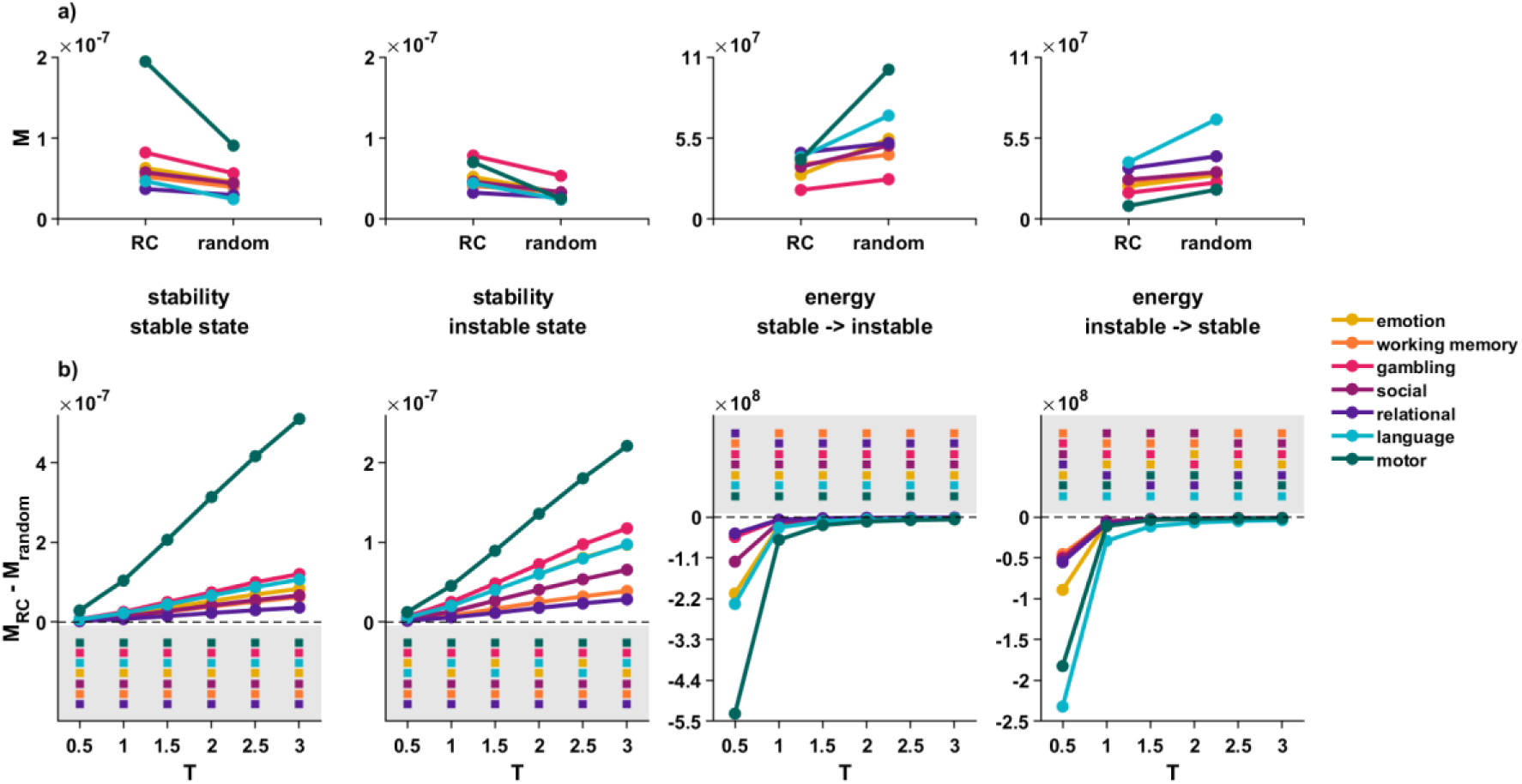
The rich club is significantly not implicated in brain state control. The rich club always contributed less to maintenance of brain states and transitions between them than a size-matched reference set of randomly chosen regions. Displayed are mean NCT values across individuals, and for each measure and task. a) A brain state was always less stable when excluding random regions from the control matrix than when excluding the rich club. Additionally, when excluding random regions, more energy was needed to traverse between brain states than when excluding the rich club. Repeated measures ANOVA found all differences to be significant for all tasks, with all *p* < 0.01. b) The direction of this effect remained the same when varying the length of the time horizon T over which trajectories were allowed to evolve; for stability, the difference between excluding the rich club vs. random regions became even more pronounced for higher T. For energy, the difference became less pronounced; importantly, the direction of the effect remained the same and the difference remained significant except for one state transition within the relational task and low T. *M* = mean value across participants, *T* = parameter choice for time horizon T

### Findings remain significant for NOS-based and partly for binary connectome weighting schemes, and across various settings of time horizon parameter

The optimal control framework within NCT evaluates brain state transitions and maintenance over time, specified by the parameter T (time horizon). To ensure the robustness of our findings regardless of the chosen time horizon, we systematically varied the parameter and re-conducted our analysis. Our main finding that excluding rich club regions versus excluding random regions had significant effect on control metrics remained consistent across all time horizons except for one state transition within the relational task when T was set to a small value (all other p < 0.001, Supplemental Tables S5 to S8). Notably, control energy estimates were less sensitive to the removal of rich club regions compared to random regions for longer time horizons, while stability estimates were more sensitive to rich club removal (see Figure 4b).

Furthermore, we explored the impact of different connectome weighting approaches by comparing our original results based on FA-weighted connectomes with results obtained with connection weights according to the number of streamlines (NOS) as well as with a binary, unweighted network. The results remained significant for the NOS-based connectome (all p < 0.05, Supplemental Tables S9 and S10). When employing a binary matrix, however, results became less consistent: We observed significant main effects for all language and motor task measures (all p < 0.001), but in the other five tasks, all energy and most stability measures (save for the gambling task and the instable states of the emotion and social task) failed to reach significance (refer to Supplemental Tables S13 and S14 and Supplemental Figures 3 and 4).

### Findings remain significant for subject-level selections of rich club members and for more closely matching the set of random regions to the rich club regarding sum of connections and similarity of connectivity profiles

To account for the considerable interindividual variability in which regions were assigned rich club status on an individual level, we also repeated the control matrix exclusions of the set of rich club regions using subject-specific rich club definitions: For each participant, we removed the top 21 highest degree nodes from the matrix of control nodes and calculated corresponding stability and control energy values. Despite the considerable interindividual variability in rich club definitions, all differences relative to a random reference set remained significant except for one state transition within the relational task (see Figure 5a, and Supplemental Tables S15 and S16).

**Figure 5.**
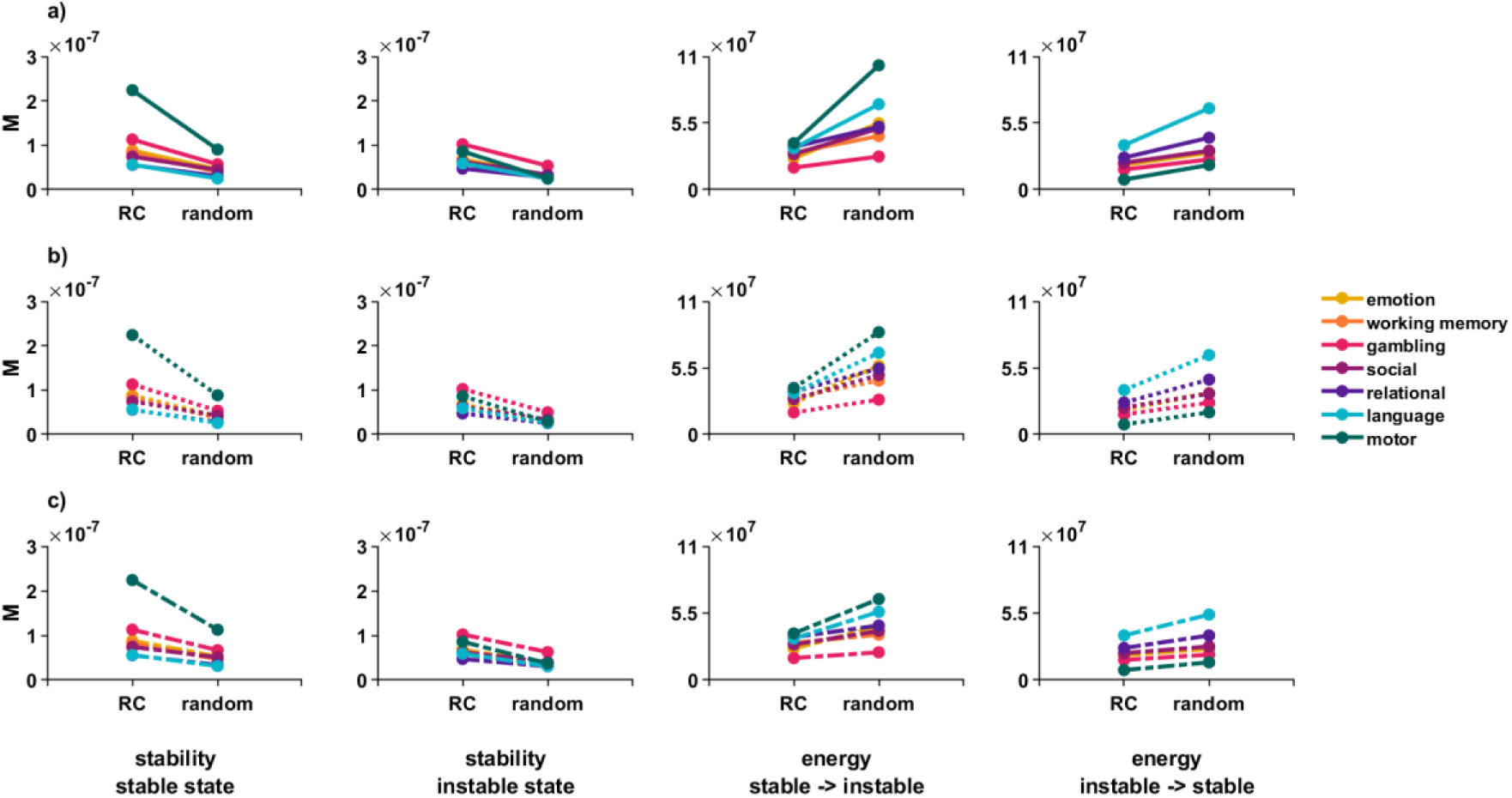
The rich-club does not control state transitions when individual variation in membership and different definitions of the reference set are taken into account. We have reported considerable individual variation in rich club members when the rich club was defined on a subject-specific level (see Figure 3). We thus repeated analyses, selecting the 21 rich club regions individually for each subject and re-calculating the impact of prohibiting those regions from exerting control versus prohibiting a size-matched reference set. Displayed are mean NCT values across individuals, for each measure, and for various sets of random regions. a) Solid line plots show the results of individual level analyses when the set of random regions was selected in the same manner as in the group level analyses, that is, fully based on the spin-test procedure described in the main text and methods section. We were able to fully replicate the effect found for a group-level rich club definition. b) Dotted plots show results of individual level analyses when the set of random regions was selected in a way in which the similarity of its members’ connectivity profiles most closely resembled that of the individual’s rich club. Again, we were able to fully replicate the group-level effect. c) Dash-dotted plots show results of individual level analyses when the set of random regions was selected in a way in which the sum of its connections most closely resembled that of the individual’s rich club. We were able to replicate the group-level effect for all stability measures and all but four state transitions; for these four transitions, differences between exclusion of the rich club and the reference set were non-significant. *M* = mean value across participants, RC = rich club

Rich club regions are inherently more interconnected than peripheral regions (van den Heuvel & Sporns, 2011) and their connectivity profiles are more similar to each other than those of a size-matched set of random regions. To ensure that our results were not spuriously inflated by simply removing regions with more mutual connections or more diverse connectivity profiles from the control set, we repeated the individual-level analyses. This time, however, we constrained the random selection of reference regions to match the rich club most closely regarding the similarity of their connectivity profile (Figure 5b) or the sum of their mutual connections (Figure 5c). The differences in stability and control energy remained significant in all cases where we compared the rich club to a connectivity-profile matched reference set. When comparing the rich club to a connection-sum-matched reference set, all differences remained significant for all stability measures and all but four state transitions, three of which concerned transitions from the instable to stable state (see Supplemental Tables S17 through S20)

### Findings largely remain significant for transitions between states associated with different tasks: significance and effect size of this comparison differ with choice of target state

So far, we have focused on *within*-task transitions, examining the impact of exclusions on state stabilization and state transitions within a given behavioral task. Behavioral tasks are designed to elicit similar brain states that differ primarily in a critical variable, such as working memory load or emotional valence. This design choice guarantees a high level of experimental control but may limit the generalizability and ecological validity of our findings, as the brain often transitions between more diverse activity patterns in real-life scenarios. To approximate more diverse transitions, we analyzed the most representative state of each task (e.g. high working memory load in the working memory task, or emotional valence in the emotion task) and investigated transitions *across* tasks. We found that even for between-task transitions, removing the rich club from the control set had a significantly reduced impact on brain state stability and brain state transitions compared to removing a size-matched reference set of random regions (see Figure 6). Effect sizes and significance of this finding varied by target state. Peripheral regions played a more critical role in control of transitions when the target state was the representative language or motor state (Figure 6); but were not differentially relevant to the rich club for some transitions towards the representative social, relational or working memory state (Figure 6; Supplemental Table S21). The mean effect size of transitions towards a target state correlated positively with the difference in that state’s stability when excluding the random reference set versus the rich club from the control matrix (*r* = 0.73; *p* = 0.0382, see also Figure 6b). This indicates that the peripheral regions are particularly crucial for transitions to states that are harder to stabilize. Importantly, we didn’t find significant correlations between effect size and the stability measure when prohibiting the rich club (*r* = 0.69, *p* = 0.0528) or when prohibiting the reference set (*r* = 0.59, *p* = 0.0873) from control; only the difference between these two measures was significantly related to effect size.

**Figure 6.**
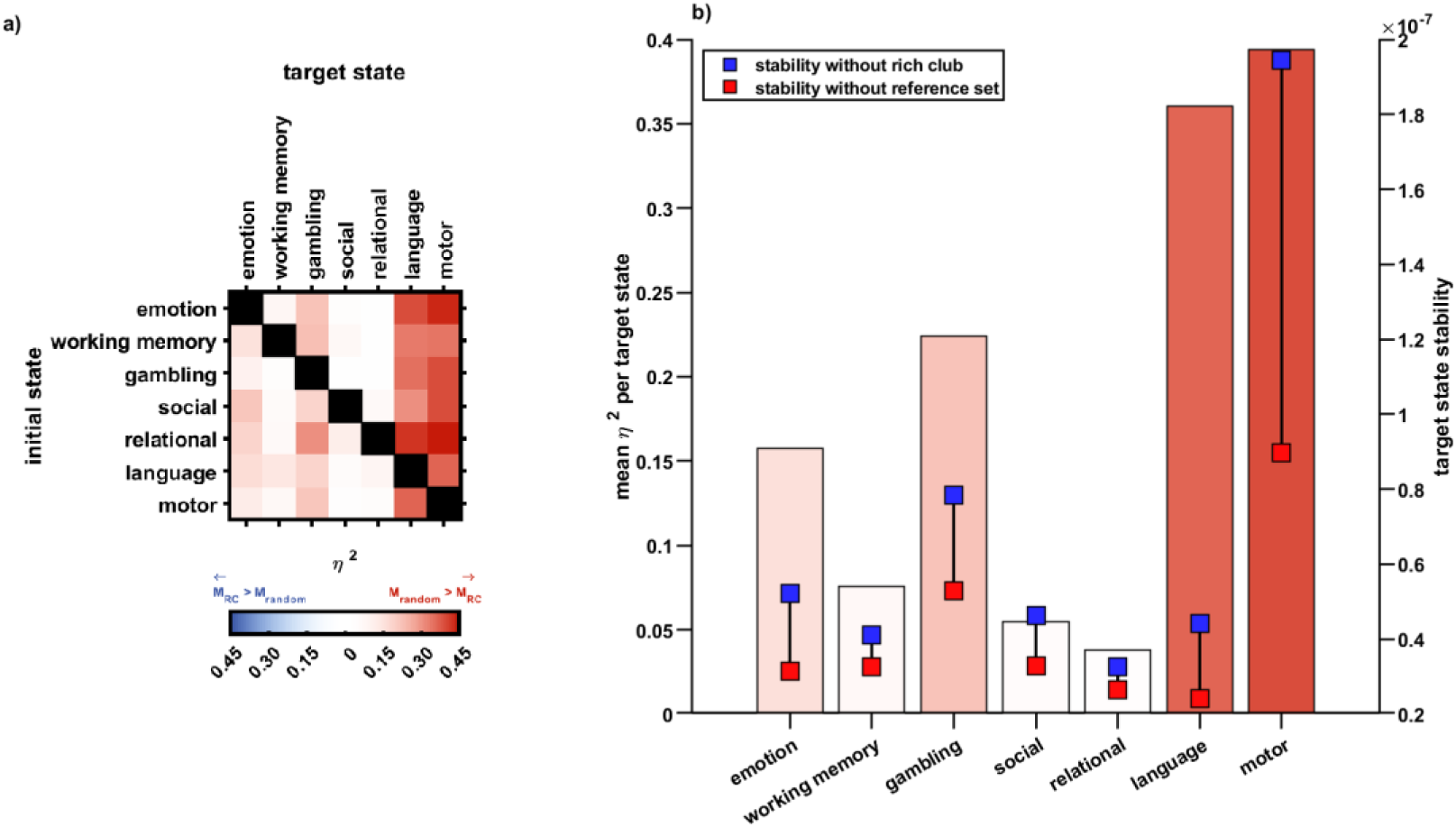
The rich club does not control brain state transitions across tasks; effect sizes depend on the target state. In reality, the brain often traverses between states that are very diverse; arguably more diverse than two states designed for one and the same experimental paradigm. We approximated this diversity by repeating our analyses for transitions across tasks, that is, from the typical brain state associated with one paradigm to that associated with a different paradigm altogether. Shown are effect sizes for the difference in control metrics when excluding the rich club vs. when excluding random regions from control. The more saturated the color, the higher the effect size. a) Repeating our findings from within-task-transitions, the periphery instead of the rich club dominated transitions across tasks: Red colors indicate a more drastic impact when excluding random regions, blue colors would indicate a more drastic impact when excluding the rich club. This effect was particularly stark when the target state to be transitioned into was associated with the language or motor task. b) The role of the periphery in controlling state transitions was particularly stark for cases in which it was, compared to the rich club, increasingly involved in stabilizing the target state: Bars show mean effect sizes off between-task-transitions per target state (left y-axis). Blue and red data points show stability of the respective state when excluding rich club regions (blue) vs. a random reference set (red) from control (right y-axis). Mean effect size was significantly related to the difference between these two stability measures, but not the respective measures themselves.

In summary, the removal of the rich club from the control set had less of an impact on brain state control than the removal of a similar group of random regions. This observation was consistent across behavioral tasks, across transitions, across rich-club definitions, across NCT parameter settings, and, to a limited degree, across connectome weighting schemes, underscoring the peripheral regions’ importance in maintaining and transitioning between diverse brain states.

### Who else, if not the rich club? Exploring the role of various cortical gradients in regional control

When comparing all regions’ mean rank of control contributions across tasks, we were able to show that high nodal degree regions in general, and the rich club in particular, contributed significantly little to optimal brain state control. In order to investigate which other features of cortical network organization might be relevant instead, we re-examined possible associations of the regional rank-measure with other candidate measures.

Previous work has highlighted the importance of peripheral over core regions for information integration and fast processing (Betzel et al., 2016; Gollo et al., 2015), for instance by proposing the concept of a “diverse club” of regions with most diverse connections across functional modules in the connectome (Bertolero et al., 2017; Lohia et al., 2024). Using the canonical 17-network-division of the cerebral cortex (Yeo et al., 2011) we calculated each region’s mean participation coefficient across participants and calculated its correlation with the regional rank measure. We found that the higher a region’s participation coefficient, the less it contributed to both the maintenance of and transitions between states (stability: *r* = 0.18; *p* = 0.0383, control energy: *r* = 0.22; *p* = 0.0164, spin test with 10,000 permutations). The similar findings for nodal degree and participation coefficient are, however, not surprising, as both measures correlated strongly at the group level (*r* = 0.54; *p* < 0.001, see also Figure 7b).

**Figure 7.**
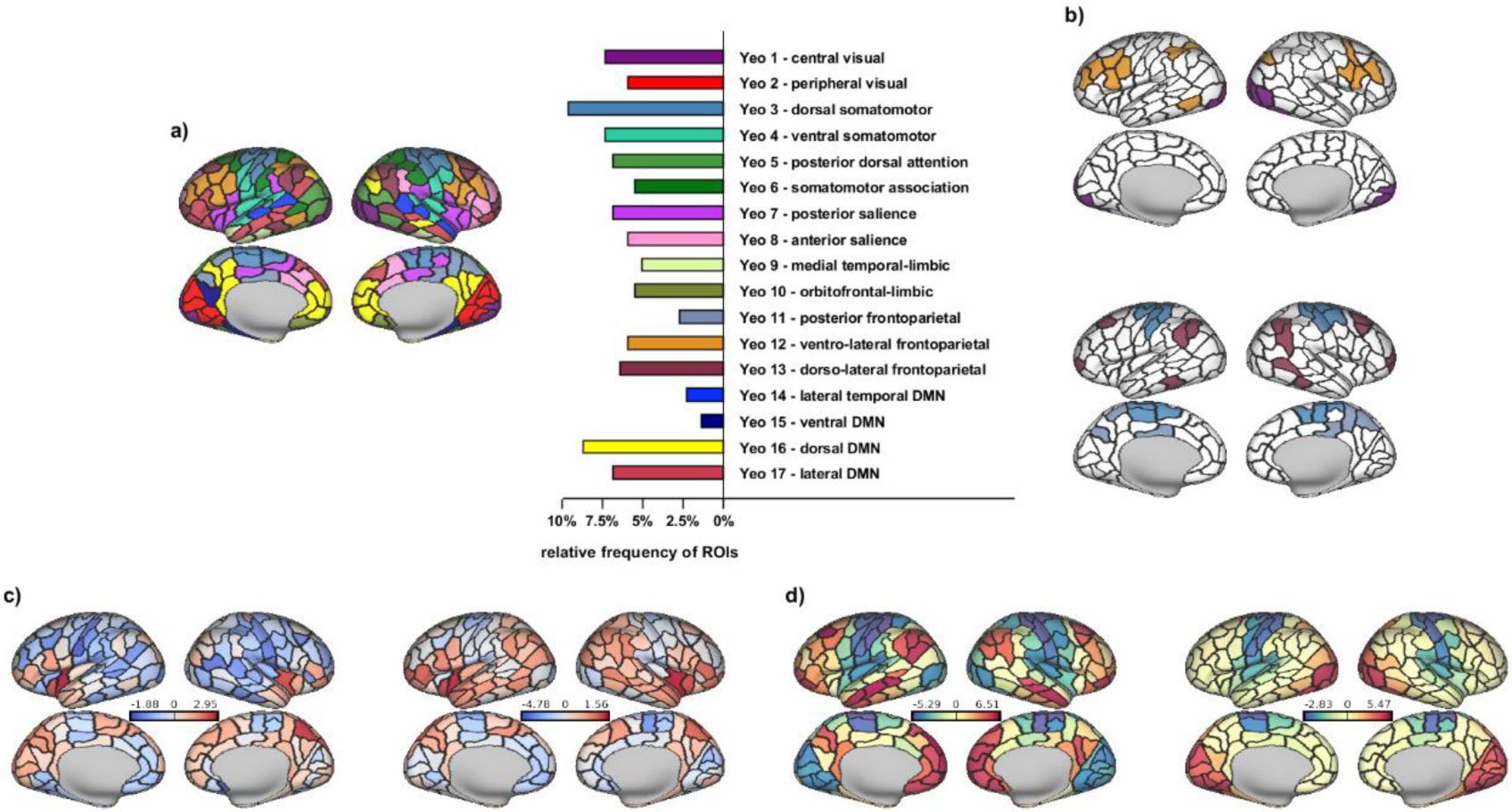
Who else, if not the rich club? Probing the role of intrinsic connectivity networks and cortical gradients in regional control. We systematically showed that the rich club does not optimally control brain state dynamics; but also that a different set of regions was task-generally implicated as control nodes (see Figure 2). Here, we further characterized this set of top control nodes by comparing it to intrinsic connectivity networks (a,b) and probing its position on various cortical gradients (c,d). a) Shows regional assignments to intrinsic connectivity networks as defined in the 17-network solution in Yeo et al. (2011); regions of interest are as defined in the Lausanne subparcellation of the Desikan-Killiany atlas. The bar chart shows the relative frequency with which a certain network was assigned to ROIs in our sample, and includes network labels. b) Regions in networks 1 and 12 contributed significantly more to regional control of brain dynamics than expected by chance (top); regions in networks 3, 11 and 13 (the latter only for stability measures) contributed significantly less than expected by chance (bottom). c) The higher the degree (*left*; z-scored mean nodal degree per region), the lower a region’s contribution to control of brain state dynamics. This mirrors our results regarding the rich club. Interestingly, we found the same relationship for a possible antagonist to degree, the participation coefficient: The higher the participation coefficient, the lower a region’s contribution to control (*right*; z-scored mean participation coefficient per region). Both measures were highly correlated on a group level in our sample. d) Regions higher on the second (*right*) principal cortical gradient as defined in Margulies et al. (2016) contributed significantly to regional control. No significant relationship was found between regional control and regional position on the first principal cortical gradient (*left*). Warm colors indicate higher positions, cool colors lower positions on the respective gradient. *DMN* = default mode network

Next, we examined the control role of various functional networks (see Supplemental Table S22, Figure 7a). We assigned every brain region to one of the 17 functional Yeo networks based on spatial overlap at the vertex-level. We then computed the mean regional rank per network, and again compared it to a spin-test based null distribution: Two networks contributed significantly more to brain state stability and brain state transitions than expected by chance (all *p* < 0.05, see Figure 7b, top): The primary visual network (Y1) and the ventrolateral division of the fronto-parietal network (Y12). Conversely, three networks contributed less to control energy than expected by chance (all *p* < 0.05), see Figure 7b, bottom: The dorsal division of the somato-motor network (Y3) and the posterior division of the fronto-parietal network (Y11) contributed less to both maintenance of states as well as transitions between states; while the dorso-lateral division of the fronto-parietal network (Y13) contributed less to maintenance of states only.

We also investigated the correlation between mean regional rank and that region’s position on the primary and secondary cortical gradient (Margulies et al., 2016). A region’s position on the secondary, but not the primary, cortical gradient was significantly negatively associated with its regional contribution to both stability (*r* = -0.47; *p* < 0.001) and control energy (*r* = -0.44; *p* < 0.001; see also Figure 7d). This suggests that regions higher along the secondary cortical gradient, which spans from motor to visual-sensory cortices, are more involved in control tasks.

Lastly, we explored possible relationships between mean rank of a region and how strongly structure and function are coupled in that region (Valk et al., 2022). We found no significant associations between mean rank regarding control contribution of a region and the extent of its structural-functional coupling.

## Discussion

The human brain’s adaptability results from its ability to stabilize behaviorally appropriate activity patterns or to flexibly adjust its state to meet current demands (Benisty et al., 2024; Braun et al., 2015; Shine et al., 2016). While the brain uses the anatomy of its white matter scaffold for these adjustments (Cornblath et al., 2020; Honey et al., 2007; Sorrentino et al., 2021), the exact mechanisms steering these transitions are not fully understood.

One candidate for general brain state control is the rich club, a group of very highly interconnected hub regions (Gollo et al., 2015; Senden et al., 2017, 2018; Towlson et al., 2013; van den Heuvel et al., 2012).

We here used optimal network control and graph-theoretical tools to investigate a possible control role of the rich club during maintenance of and transition between different brain states, constrained by seven canonical fMRI tasks. We found that rich club regions and high-degree regions in general had consistently less impact on control metrics than peripheral regions. This finding was robust across different parameters and definitions, and only vanished partially when utilizing a binary connectome map, which is a highly simplified representation of the brain’s anatomical scaffold.

Our findings challenge the traditional view that high degree regions such as the rich club form an integrative core necessary for the control of brain function (de Pasquale et al., 2016; Deco et al., 2017, 2021; Zink et al., 2021) and efficient brain state transitions (Betzel et al., 2016). Quite on the contrary, we did not observe a single brain state trajectory to which the rich club contributed more control energy than the reference set. Previous work suggests that the role of hub regions in network control may not be universal and state-general, but depends on the ease of reaching a brain state, with hub regions primarily controlling easily reachable states (Gu et al., 2015). However, in our study, we found no clear evidence that transitions to more stable, easily reachable brain states required more contributions from the rich club than transitions to less stable states. This indicates that either the brain states we examined might be too difficult to reach for the rich club’s control, or that the rich club contributes less to the control of brain state dynamics than previously thought.

Interestingly, regions with a high participation coefficient also did not emerge as top control nodes. The participation coefficient characterizes regions whose connections connect across module boundaries within the connectome. It has been argued that these diverse regions may be more involved than high-degree rich club regions in the orchestration of information exchange in the brain network (Bertolero et al., 2017; Gollo et al., 2015; Lohia et al., 2024). However, the present data suggest that the putative “diverse club” contributes as little to the control of brain state dynamics as the rich club and that indeed, increased cognitive effort might instead be associated more with diminishing importance of clubness and decreased modularity in the brain (Finc et al., 2017; Hearne et al., 2017; Kitzbichler et al., 2011).

Instead, a region’s effectiveness in controlling brain state transitions may depend more on its membership in certain intrinsic connectivity networks or its position on specific cortical gradients of cortical functional connectivity. Specifically, regions within the primary visual network (Y1) and the ventrolateral division of the fronto-parietal network (Y12) exhibited especially high task-general control influences.

Our findings demonstrate that the fronto-parietal network exhibited exceptionally high task-general control influences. This result is consistent with the dual network account of cognitive control, which identifies the fronto-parietal network as one of two principal control networks (Cornblath et al., 2020; Dosenbach et al., 2007; Gu et al., 2015; Rosen et al., 2016; W. W. Seeley et al., 2007). According to this perspective, the fronto-parietal network is implicated in domain-general cognitive control and executive functioning (Schultz et al., 2022; W. P. Seeley, 2020), particularly over shorter timescales (Power & Petersen, 2013). However, the dual network account presents a more nuanced view by differentiating between state transitions and state maintenance, attributing these functions to the fronto-parietal and cingulo-opercular networks, respectively (Dosenbach et al., 2007). Interestingly, our study revealed that the ventro-lateral portion of the fronto-parietal network was task-generally involved in both the transitions between and maintenance of brain states. This suggests that the fronto-parietal network’s role in cognitive control may be more integrative than previously understood, encompassing both dynamic adjustments and sustained control processes. It is also noteworthy that in this present study, the posterior section, and for stability measures also the dorso-lateral section, of the fronto-parietal network had much like the rich club, a significantly negligible contribution to control of brain states. This finding underscores the heterogeneity within the fronto-parietal network itself, challenging the idea that the entire fronto-parietal network contributes to brain control.

The participation of the primary visual network (Y1) in control processes was also reflected in our findings regarding primary and secondary cortical gradients: Being positioned higher on the secondary gradient distinguishing somatomotor-regions (lower) from visual regions (higher; Margulies et al., 2016) was significantly related to high task-general control participation. The striate cortex might not only hold a special position as the starting point of the cortical hierarchy (Mesulam, 1998), but also as starting point for higher cognitive functions such as cognitive control (Gavornik & Bear, 2014; Muckli, 2010). Activity in occipital cortex, particularly in the calcarine cortex and area V1, has been shown to precede cognitive control failures (Cieslik et al., 2024; Hamm et al., 2012; Su et al., 2020) and to encode multisensory integration (Murray et al., 2016). The visual network has also been shown to merge together with a frontoparietal module under increasing cognitive complexity of a task (Hearne et al., 2017). The classical account posits the primary visual network as a specific unisensory processing area, though this, also in light of our findings here, might constitute a significant oversimplification (Muckli, 2010).

Methodological considerations need to be made: Following previous research (Braun et al., 2021) we use first-level analysis results - specifically, spatial patterns of beta weights - as the basis for brain-state definition. Therefore, brain states are resulting from statistical maps rather than direct BOLD-level activity patterns. Utilizing these beta weighted maps enabled us to base brain states on behaviorally constrained task fMRI data in the attempt for greater ecological validity. This approach is different from previous research that has capitalized on either simulated or resting state data (Betzel et al., 2016; Gu et al., 2015, 2017). It should be noted that when using such beta-weighted maps as the basis for brain states, nodes similarly active during both initial and target states are not discernable from those inactive in both states. The inability to discriminate between task-generally very active nodes and task-generally very inactive nodes might be a possible confound in resulting control metrics. Furthermore, the specific and circumscribed nature of the task data collected within the Human Connectome Project should be considered when interpreting our results.

Given the ongoing debate about the precise interpretation of modal and average controllability metrics (Pasqualetti et al., 2019; Suweis et al., 2019; Tu et al., 2018), we employed a third methodological approach in our choice of optimal control analyses. Nonetheless, replicating our results using average and modal controllability metrics might prove fruitful. These two approaches allow for distinct investigations of easily reachable and difficult to reach brain states (Karrer et al., 2020). Similar to Luppi et al. (2023) we found that the effect size of the control impact difference between the rich club and the reference set was partially related to target state characteristics. Other research has linked cognitive load to the specifics of state trajectories (Cornblath et al., 2020). Systematically considering target state specifics, such as associated cognitive load or reachability (Gu et al., 2015) can further elucidate network control processes further and enhance approaches that respects current methodological boundaries of network control theoretical analyses (see also Karrer et al., 2020; Pasqualetti et al., 2019; Tu et al., 2018). Combined with the use of time-resolved activity patterns, such as source-reconstructed EEG or MEG data, this would allow for a closer study of the dynamics of control, including possibly regionally different timescales (Gollo et al., 2015)

## Methods

### Dataset

Data used here was sourced exclusively from the Human Connectome Project, specifically the subset of data from 100 unrelated subjects. The HCP provides data from seven tasks, an emotion matching task, a version of an n-back working memory task, a gambling task, a language task, a motor task eliciting movement, a relational task requiring participants to match visual properties of objects, and a social task assessing whether participants think of objects as having a random or social interaction. We used preprocessed (Glasser et al., 2013; Van Essen et al., 2013) first-level contrasts from all seven tasks (see Figure 1). See Barch et al. (2013) for an overview over the HCP’s choice of task paradigms.

Task data was combined with preprocessed diffusion-weighted MRI data as provided by the HCP (see Van Essen et al., 2013 regarding acquisition pipeline) to reconstruct subject-specific connectomes.

### Connectome Reconstruction

We processed the structural data using the connectivity reconstruction toolbox CATO (de Lange et al., 2023), including application of CATO’s minimal preprocessing pipeline to create cortical parcellations from the Freesurfer output. Cortical parcellations were created based on the Lausanne sub-parcellation of the Desikan-Killiany atlas (Cammoun et al., 2012), resulting in 219 cortical regions. White matter fiber organization was estimated using a combined approach of diffusion tensor imaging (DTI) and generalized Q-sampling imaging (GQI), leveraging the strengths of both methods. DTI modeling was here performed in voxels with one peak, GQI in those with multiple peaks (de Lange et al., 2023). Fiber tracking in CATO is based on a version of the deterministic “Fiber Assignment by Continuous Tracking” (FACT) algorithm originally described by (Mori et al., 1999); see de Lange et al. (2023) for details regarding the procedure. For any two regions in the parcellation, we determined the shortest fiber segment (if any) and calculated the mean FA of all voxels in each fiber segment, weighted by the length of the traversed path through each voxel. This information was then used to reconstruct weighted structural networks for each participant. While FA-weighted connectomes were used throughout the main analysis, we additionally derived unweighted connectomes and connectomes weighted by the number of streamlines for parameter exploration. Of the initial *N* = 100, *N* = 98 participants (*n* = 45 male, *n* = 53 female; mode of age = 33) were included in our investigation: Following the procedure described in van den Heuvel & Sporns (2019), we tested for outliers based on a group average prevalence matrix characterizing the average number of times a connection between two nodes was observed across participants. We excluded data of two participants whose connectome maps significantly deviated from the group average to satisfy criteria for removal. All analyses used the full set of N = 98 participants, except for the exploratory analyses with NOS-weighted connectomes, where one additional participant with outlier values was excluded (see Supplemental Figure 3), resulting in a set of N = 97 for these analyses.

### Brain state definition

We based brain states on the two main experimental contrasts for all tasks; regarding the motor task, we opted for those contrasts that contained data from both hemispheres. Individual cortical parcellations were brought into CIFTI format using Freesurfer and connectome workbench (Marcus et al., 2011), and then used to parcellate functional data, allowing us to respect individual differences in regional boundaries. We defined regional activity values by calculating mean activation over time and per region, resulting in two Lausanne-parcellated brain states per task and participant. Visualizations using brain projections in this study are always created using connectome workbench.

### Definition of the rich club and the null model

We established a rich club regime following the procedure detailed in Riedel et al. (2022). Using the Brain Connectivity Toolbox (Rubinov & Sporns, 2010; version 2019_03_03), we first computed weighted rich club coefficients for our empirical connectomes (rich_club_wu.m), and then for 10,000 random brain networks, each connection randomly rewired 10 times while preserving degree distribution (randmio_und.m). The normalized rich club coefficient *Φ* for each level *k* of the degree distribution is the ratio of the empirical coefficient over the random coefficient, averaged across all 10,000 iterations. Setting a cut-off at *k_max(Φ)_* (see Figure 3b), on average 9.69% of an individual’s nodes (around 21 nodes) were considered rich club members. Group-level rich club definition was based on the 9,69% top ranking nodes across subjects, that is, those 9.69% of nodes that qualified most consistently as rich club members across participants, resulting in 21 regions constituting the group-level rich club (see Figure 3a; c-d). Individually specific rich clubs were subsequently based on the 21 top degree nodes of an individual, ensuring that while the exact composition of the rich club was subject-specific, the number of rich club members remained matched across individuals.

For inferential evaluations, we compared results of excluding those 21 rich club regions from the control matrix to a null model in which we excluded 21 semi-randomly selected regions instead. Our null models are based on a spin model paradigm (Alexander-Bloch et al., 2018; Váša et al., 2018; code available under https://github.com/frantisekvasa/rotate_parcellation) which randomly rotates spherical surface projections of the atlas while preserving hemispheric symmetry and contiguity. For NCT analyses, this procedure was repeated 50 times, resulting in 50 rotated brain maps. The mean control energy and stability of excluding 21 regions across these 50 null model maps were statistically compared to the results of excluding the rich club from the matrix of control nodes.

We also use the spin-test model to evaluate correlations of various region specific properties (nodal degree, participation coefficient, membership in functional networks, as well as the position on cortical gradients) and regional control energy. The spin model permuted values of interest 10,000 times to construct null distributions against which the empirical values were compared to determine statistical significance of the correlations.

### Control energy and stability

The control energy framework employed here is a linear approximation of brain dynamics based on a) a stabilized structural connectivity matrix *S* of a subject, b) two vectors *x(t)* describing brain states at a given point in time, c) a control matrix *C* delineating which regions are allowed to control brain dynamics, and d) a vector of control energy *u(t)* representing the control input into the control nodes at a given point in time (Braun et al., 2021; see also Betzel et al., 2016; Karrer et al., 2020):

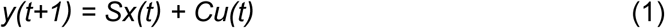

This formula can be solved for an optimized value of parameter *u* minimizing both required control energy as well as length of trajectory between the initial and target state through state space (Braun et al., 2021; Karrer et al., 2020). We define optimal control energyfor the transition between two distinct brain states as squared integral of a region’s energy contribution over time, averaged across all 219 brain regions. We define stability of a state accordingly, as the optimal control effort needed for maintenance of a state, i.e as the inverse of the squared integral of a region’s energy contribution over time, averaged across all 219 brain regions, when defining the same state as initial and target state (Braun et al., 2021). All NCT analyses were implemented in MATLAB R2021b, based on NCT-functions publicly provided by Braun et al. (2021; especially optim_fun.m, available under https://github.com/ursbraun/network_control_and_dopamine). Within our main analyses, we set the parameters for time horizon *T* = 1 and penalization parameter *ρ* = 1; however, we also systematically investigate the impact of changing the values for time horizon *T*. We transformed raw NCT output by dividing by the number of timesteps given by our time horizon T, allowing for comparisons across different settings of the time parameter.

The control energy framework was applied with three distinct manipulations of the control matrix C. In order to probe the general optimal control energy and stability, we included all 219 regions into the control set, enabling all regions to contribute to state transitions and maintenance. To probe the role of the rich club, we utilized a reduced set of control nodes We either excluded rich club regions from the control matrix by setting their respective entries in the control matrix to zero, or excluded a size-matched set of random regions from the control set. In order to probe individual contributions of brain regions, we removed single brain regions from the control set and evaluated the change in control energy and stability against the full control set. Please note that removing nodes from the control matrix does not remove them from the connectome; they can still be influenced and controlled by other nodes, but cannot exert control themselves (Betzel et al., 2016).

### Statistics

Repeated measures ANOVA was used to statistically compare NCT values (optimal control energy and stability) between the two states assigned to a task, as well as to study the effect of prohibiting control of the rich club vs. random regions. In line with previous research (Braun et al., 2021) we controlled for mean nodal activity difference between states A and B to ensure that possible differences in control metrics were not merely reflecting differences in brain activation. Mean activation per ROI for each participant and brain state was extracted, and the difference in mean regional activation between states A and B was included in the repeated measures model as a covariate. Post hoc paired sample t tests were conducted to confirm direction of significant main effects.

For regional NCT metrics and their association with regionally specific properties, we used product-moment correlation coefficients to quantify this relationship. Regions were rank-ordered according to their impact on stability and control energy metrics, and each region’s mean rank across tasks and across stability and energy measures was correlated with the regional property of interest (specifically, the mean of that variable per node and across participants). Correlations were evaluated against a null distribution of values obtained by permuting regional rank 10,000 times according to the spin test model, calculating how often the empirical value was exceeded by permuted values, forming the basis for the assigned p-values.

## References

Alexander-Bloch, A. F., Shou, H., Liu, S., Satterthwaite, T. D., Glahn, D. C., Shinohara, R. T., Vandekar, S. N., & Raznahan, A. (2018). On testing for spatial correspondence between maps of human brain structure and function. NeuroImage, 178, 540–551. 10.1016/j.neuroimage.2018.05.070

Alstott, J., Panzarasa, P., Rubinov, M., Bullmore, E. T., & Vértes, P. E. (2014). A Unifying Framework for Measuring Weighted Rich Clubs. Scientific Reports, 4(1), 7258. 10.1038/srep07258

Barch, D. M., Burgess, G. C., Harms, M. P., Petersen, S. E., Schlaggar, B. L., Corbetta, M., Glasser, M. F., Curtiss, S., Dixit, S., Feldt, C., Nolan, D., Bryant, E., Hartley, T., Footer, O., Bjork, J. M., Poldrack, R., Smith, S., Johansen-Berg, H., Snyder, A. Z., & Van Essen, D. C. (2013). Function in the human connectome: Task-fMRI and individual differences in behavior. NeuroImage, 80, 169–189. 10.1016/j.neuroimage.2013.05.033

Bassett, D. S., Wymbs, N. F., Rombach, M. P., Porter, M. A., Mucha, P. J., & Grafton, S. T. (2013). Task-Based Core-Periphery Organization of Human Brain Dynamics. PLoS Computational Biology, 9(9), e1003171. 10.1371/journal.pcbi.1003171

Benisty, H., Barson, D., Moberly, A. H., Lohani, S., Tang, L., Coifman, R. R., Crair, M. C., Mishne, G., Cardin, J. A., & Higley, M. J. (2024). Rapid fluctuations in functional connectivity of cortical networks encode spontaneous behavior. Nature Neuroscience, 27(1), 148–158. 10.1038/s41593-023-01498-y

Bertolero, M. A., Yeo, B. T. T., & D’Esposito, M. (2017). The diverse club. Nature Communications, 8(1), 1277. 10.1038/s41467-017-01189-w

Betzel, R. F., Gu, S., Medaglia, J. D., Pasqualetti, F., & Bassett, D. S. (2016). Optimally controlling the human connectome: The role of network topology. Scientific Reports, 6(1), 30770. 10.1038/srep30770

Braun, U., Harneit, A., Pergola, G., Menara, T., Schäfer, A., Betzel, R. F., Zang, Z., Schweiger, J. I., Zhang, X., Schwarz, K., Chen, J., Blasi, G., Bertolino, A., Durstewitz, D., Pasqualetti, F., Schwarz, E., Meyer-Lindenberg, A., Bassett, D. S., & Tost, H. (2021). Brain network dynamics during working memory are modulated by dopamine and diminished in schizophrenia. Nature Communications, 12(1), 3478. 10.1038/s41467-021-23694-9

Braun, U., Schäfer, A., Walter, H., Erk, S., Romanczuk-Seiferth, N., Haddad, L., Schweiger, J. I., Grimm, O., Heinz, A., Tost, H., Meyer-Lindenberg, A., & Bassett, D. S. (2015). Dynamic reconfiguration of frontal brain networks during executive cognition in humans. Proceedings of the National Academy of Sciences, 112(37), 11678–11683. 10.1073/pnas.1422487112

Cammoun, L., Gigandet, X., Meskaldji, D., Thiran, J. P., Sporns, O., Do, K. Q., Maeder, P., Meuli, R., & Hagmann, P. (2012). Mapping the human connectome at multiple scales with diffusion spectrum MRI. Journal of Neuroscience Methods, 203(2), 386–397. 10.1016/j.jneumeth.2011.09.031

Cieslik, E. C., Ullsperger, M., Gell, M., Eickhoff, S. B., & Langner, R. (2024). Success versus failure in cognitive control: Meta-analytic evidence from neuroimaging studies on error processing. Neuroscience & Biobehavioral Reviews, 156, 105468. 10.1016/j.neubiorev.2023.105468

Cole, M. W., Ito, T., Bassett, D. S., & Schultz, D. H. (2016). Activity flow over resting-state networks shapes cognitive task activations. Nature Neuroscience, 19(12), 1718–1726. 10.1038/nn.4406

Collin, G., Sporns, O., Mandl, R. C. W., & van den Heuvel, M. P. (2014). Structural and Functional Aspects Relating to Cost and Benefit of Rich Club Organization in the Human Cerebral Cortex. Cerebral Cortex, 24(9), 2258–2267. 10.1093/cercor/bht064

Cornblath, E. J., Ashourvan, A., Kim, J. Z., Betzel, R. F., Ciric, R., Adebimpe, A., Baum, G. L., He, X., Ruparel, K., Moore, T. M., Gur, R. C., Gur, R. E., Shinohara, R. T., Roalf, D. R., Satterthwaite, T. D., & Bassett, D. S. (2020). Temporal sequences of brain activity at rest are constrained by white matter structure and modulated by cognitive demands. Communications Biology, 3(1), 261. 10.1038/s42003-020-0961-x

de Lange, S. C., Helwegen, K., & van den Heuvel, M. P. (2023). Structural and functional connectivity reconstruction with CATO - A Connectivity Analysis TOolbox. NeuroImage, 273, 120108. 10.1016/j.neuroimage.2023.120108

de Pasquale, F., Della Penna, S., Sporns, O., Romani, G. L., & Corbetta, M. (2016). A Dynamic Core Network and Global Efficiency in the Resting Human Brain. Cerebral Cortex, 26(10), 4015–4033. 10.1093/cercor/bhv185

de Reus, M. A., & van den Heuvel, M. P. (2014). Simulated rich club lesioning in brain networks: A scaffold for communication and integration? Frontiers in Human Neuroscience, 8. 10.3389/fnhum.2014.00647

Deco, G., Kringelbach, M. L., Jirsa, V. K., & Ritter, P. (2017). The dynamics of resting fluctuations in the brain: Metastability and its dynamical cortical core. Scientific Reports, 7(1), 3095. 10.1038/s41598-017-03073-5

Deco, G., Vidaurre, D., & Kringelbach, M. L. (2021). Revisiting the global workspace orchestrating the hierarchical organization of the human brain. Nature Human Behaviour, 5(4), 497–511. 10.1038/s41562-020-01003-6

Dimulescu, C., Gareayaghi, S., Kamp, F., Fromm, S., Obermayer, K., & Metzner, C. (2021). Structural Differences Between Healthy Subjects and Patients With Schizophrenia or Schizoaffective Disorder: A Graph and Control Theoretical Perspective. Frontiers in Psychiatry, 12, 669783. 10.3389/fpsyt.2021.669783

Dosenbach, N. U. F., Fair, D. A., Miezin, F. M., Cohen, A. L., Wenger, K. K., Dosenbach, R. A. T., Fox, M. D., Snyder, A. Z., Vincent, J. L., Raichle, M. E., Schlaggar, B. L., & Petersen, S. E. (2007). Distinct brain networks for adaptive and stable task control in humans. Proceedings of the National Academy of Sciences, 104(26), 11073–11078. 10.1073/pnas.0704320104

Finc, K., Bonna, K., Lewandowska, M., Wolak, T., Nikadon, J., Dreszer, J., Duch, W., & Kühn, S. (2017). Transition of the functional brain network related to increasing cognitive demands. Human Brain Mapping, 38(7), 3659–3674. 10.1002/hbm.23621

Gavornik, J. P., & Bear, M. F. (2014). Higher brain functions served by the lowly rodent primary visual cortex. Learning & Memory, 21(10), 527–533. 10.1101/lm.034355.114

Glasser, M. F., Sotiropoulos, S. N., Wilson, J. A., Coalson, T. S., Fischl, B., Andersson, J. L., Xu, J., Jbabdi, S., Webster, M., Polimeni, J. R., Van Essen, D. C., & Jenkinson, M. (2013). The minimal preprocessing pipelines for the Human Connectome Project. NeuroImage, 80, 105–124. 10.1016/j.neuroimage.2013.04.127

Gollo, L. L., Zalesky, A., Hutchison, R. M., van den Heuvel, M., & Breakspear, M. (2015). Dwelling quietly in the rich club: Brain network determinants of slow cortical fluctuations. Philosophical Transactions of the Royal Society B: Biological Sciences, 370(1668), 20140165. 10.1098/rstb.2014.0165

Golos, M., Jirsa, V., & Daucé, E. (2015). Multistability in Large Scale Models of Brain Activity. PLOS Computational Biology, 11(12), e1004644. 10.1371/journal.pcbi.1004644

Gonzalez-Castillo, J., & Bandettini, P. A. (2018). Task-based dynamic functional connectivity: Recent findings and open questions. NeuroImage, 180, 526–533. 10.1016/j.neuroimage.2017.08.006

Greene, A. S., Horien, C., Barson, D., Scheinost, D., & Constable, R. T. (2023). Why is everyone talking about brain state? Trends in Neurosciences, 46(7), 508–524. 10.1016/j.tins.2023.04.001

Griffa, A., & Van den Heuvel, M. P. (2018). Rich-club neurocircuitry: Function, evolution, and vulnerability. Dialogues in Clinical Neuroscience, 20(2), 121–132. 10.31887/DCNS.2018.20.2/agriffa

Gu, S., Betzel, R. F., Mattar, M. G., Cieslak, M., Delio, P. R., Grafton, S. T., Pasqualetti, F., & Bassett, D. S. (2017). Optimal trajectories of brain state transitions. NeuroImage, 148, 305–317. 10.1016/j.neuroimage.2017.01.003

Gu, S., Pasqualetti, F., Cieslak, M., Telesford, Q. K., Yu, A. B., Kahn, A. E., Medaglia, J. D., Vettel, J. M., Miller, M. B., Grafton, S. T., & Bassett, D. S. (2015). Controllability of structural brain networks. Nature Communications, 6(1), 8414. 10.1038/ncomms9414

Hagmann, P., Cammoun, L., Gigandet, X., Meuli, R., Honey, C. J., Wedeen, V. J., & Sporns, O. (2008). Mapping the Structural Core of Human Cerebral Cortex. PLoS Biology, 6(7), e159. 10.1371/journal.pbio.0060159

Hahn, T., Winter, N. R., Ernsting, J., Gruber, M., Mauritz, M. J., Fisch, L., Leenings, R., Sarink, K., Blanke, J., Holstein, V., Emden, D., Beisemann, M., Opel, N., Grotegerd, D., Meinert, S., Heindel, W., Witt, S., Rietschel, M., Nöthen, M. M., … Repple, J. (2023). Genetic, individual, and familial risk correlates of brain network controllability in major depressive disorder. Molecular Psychiatry, 28(3), 1057–1063. 10.1038/s41380-022-01936-6

Hamm, J. P., Dyckman, K. A., McDowell, J. E., & Clementz, B. A. (2012). Pre-Cue Fronto-Occipital Alpha Phase and Distributed Cortical Oscillations Predict Failures of Cognitive Control. The Journal of Neuroscience, 32(20), 7034–7041. 10.1523/JNEUROSCI.5198-11.2012

Hearne, L. J., Cocchi, L., Zalesky, A., & Mattingley, J. B. (2017). Reconfiguration of Brain Network Architectures between Resting-State and Complexity-Dependent Cognitive Reasoning. The Journal of Neuroscience, 37(35), 8399–8411. 10.1523/JNEUROSCI.0485-17.2017

Honey, C. J., Kötter, R., Breakspear, M., & Sporns, O. (2007). Network structure of cerebral cortex shapes functional connectivity on multiple time scales. Proceedings of the National Academy of Sciences, 104(24), 10240–10245. 10.1073/pnas.0701519104

Jeganathan, J., Perry, A., Bassett, D. S., Roberts, G., Mitchell, P. B., & Breakspear, M. (2018). Fronto-limbic dysconnectivity leads to impaired brain network controllability in young people with bipolar disorder and those at high genetic risk. NeuroImage: Clinical, 19, 71–81. 10.1016/j.nicl.2018.03.032

Jenkinson, M., Beckmann, C. F., Behrens, T. E. J., Woolrich, M. W., & Smith, S. M. (2012). FSL. NeuroImage, 62(2), 782–790. 10.1016/j.neuroimage.2011.09.015

Karrer, T. M., Kim, J. Z., Stiso, J., Kahn, A. E., Pasqualetti, F., Habel, U., & Bassett, D. S. (2020). A practical guide to methodological considerations in the controllability of structural brain networks. Journal of Neural Engineering, 17(2), 026031. 10.1088/1741-2552/ab6e8b

Kim, J. Z., Soffer, J. M., Kahn, A. E., Vettel, J. M., Pasqualetti, F., & Bassett, D. S. (2018). Role of graph architecture in controlling dynamical networks with applications to neural systems. Nature Physics, 14(1), 91–98. 10.1038/nphys4268

Kitzbichler, M. G., Henson, R. N. A., Smith, M. L., Nathan, P. J., & Bullmore, E. T. (2011). Cognitive Effort Drives Workspace Configuration of Human Brain Functional Networks. The Journal of Neuroscience, 31(22), 8259–8270. 10.1523/JNEUROSCI.0440-11.2011

Lohia, K., Soans, R. S., Saxena, R., Mahajan, K., & Gandhi, T. K. (2024). Distinct rich and diverse clubs regulate coarse and fine binocular disparity processing: Evidence from stereoscopic task-based fMRI. IScience, 109831. 10.1016/j.isci.2024.109831

Luppi, A. I., Singleton, S. P., Hansen, J. Y., Bzdok, D., Kuceyeski, A., Betzel, R. F., & Misic, B. (2023). *Transitions between cognitive topographies: Contributions of network structure, neuromodulation, and disease* [Preprint]. Neuroscience. 10.1101/2023.03.16.532981

Marcus, D. S., Harwell, J., Olsen, T., Hodge, M., Glasser, M. F., Prior, F., Jenkinson, M., Laumann, T., Curtiss, S. W., & Van Essen, D. C. (2011). Informatics and Data Mining Tools and Strategies for the Human Connectome Project. Frontiers in Neuroinformatics, 5. 10.3389/fninf.2011.00004

Margulies, D. S., Ghosh, S. S., Goulas, A., Falkiewicz, M., Huntenburg, J. M., Langs, G., Bezgin, G., Eickhoff, S. B., Castellanos, F. X., Petrides, M., Jefferies, E., & Smallwood, J. (2016). Situating the default-mode network along a principal gradient of macroscale cortical organization. Proceedings of the National Academy of Sciences, 113(44), 12574–12579. 10.1073/pnas.1608282113

Medaglia, J. D. (2019). Clarifying cognitive control and the controllable connectome. WIREs Cognitive Science, 10(1), e1471. 10.1002/wcs.1471

Mesulam, M. (1998). From sensation to cognition. Brain, 121(6), 1013–1052. 10.1093/brain/121.6.1013

Mori, S., Crain, B. J., Chacko, V. P., & Van Zijl, P. C. M. (1999). Three-dimensional tracking of axonal projections in the brain by magnetic resonance imaging. Annals of Neurology, 45(2), 265–269. 10.1002/1531-8249(199902)45:2%3C265::aid-ana21%3E3.0.co;2-3

Muckli, L. (2010). What are we missing here? Brain imaging evidence for higher cognitive functions in primary visual cortex V1. International Journal of Imaging Systems and Technology, 20(2), 131–139. 10.1002/ima.20236

Murray, M. M., Thelen, A., Thut, G., Romei, V., Martuzzi, R., & Matusz, P. J. (2016). The multisensory function of the human primary visual cortex. Neuropsychologia, 83, 161–169. 10.1016/j.neuropsychologia.2015.08.011

Parkes, L., Kim, J. Z., Stiso, J., Brynildsen, J. K., Cieslak, M., Covitz, S., Gur, R. E., Gur, R. C., Pasqualetti, F., Shinohara, R. T., Zhou, D., Satterthwaite, T. D., & Bassett, D. S. (2024). A network control theory pipeline for studying the dynamics of the structural connectome. Nature Protocols. 10.1038/s41596-024-01023-w

Pasqualetti, F., Gu, S., & Bassett, D. S. (2019). RE: Warnings and caveats in brain controllability. NeuroImage, 197, 586–588. 10.1016/j.neuroimage.2019.05.001

Power, J. D., & Petersen, S. E. (2013). Control-related systems in the human brain. Current Opinion in Neurobiology, 23(2), 223–228. 10.1016/j.conb.2012.12.009

Riedel, L., van den Heuvel, M. P., & Markett, S. (2022). Trajectory of rich club properties in structural brain networks. Human Brain Mapping, 43(14), 4239–4253. 10.1002/hbm.25950

Rosen, M. L., Stern, C. E., Michalka, S. W., Devaney, K. J., & Somers, D. C. (2016). Cognitive Control Network Contributions to Memory-Guided Visual Attention. Cerebral Cortex, 26(5), 2059–2073. 10.1093/cercor/bhv028

Rubinov, M. (2016). Constraints and spandrels of interareal connectomes. Nature Communications, 7(1), 13812. 10.1038/ncomms13812

Rubinov, M., & Sporns, O. (2010). Complex network measures of brain connectivity: Uses and interpretations. NeuroImage, 52(3), 1059–1069. 10.1016/j.neuroimage.2009.10.003

Schultz, D. H., Ito, T., & Cole, M. W. (2022). Global connectivity fingerprints predict the domain generality of multiple-demand regions. Cerebral Cortex, 32(20), 4464–4479. 10.1093/cercor/bhab495

Seeley, W. P. (2020). Attentional engines: A perceptual theory of the arts. Oxford University Press.

Seeley, W. W., Menon, V., Schatzberg, A. F., Keller, J., Glover, G. H., Kenna, H., Reiss, A. L., & Greicius, M. D. (2007). Dissociable Intrinsic Connectivity Networks for Salience Processing and Executive Control. The Journal of Neuroscience, 27(9), 2349–2356. 10.1523/JNEUROSCI.5587-06.2007

Senden, M., Deco, G., de Reus, M. A., Goebel, R., & van den Heuvel, M. P. (2014). Rich club organization supports a diverse set of functional network configurations. NeuroImage, 96, 174–182. 10.1016/j.neuroimage.2014.03.066

Senden, M., Reuter, N., van den Heuvel, M. P., Goebel, R., & Deco, G. (2017). Cortical rich club regions can organize state-dependent functional network formation by engaging in oscillatory behavior. NeuroImage, 146, 561–574. 10.1016/j.neuroimage.2016.10.044

Senden, M., Reuter, N., van den Heuvel, M. P., Goebel, R., Deco, G., & Gilson, M. (2018). Task-related effective connectivity reveals that the cortical rich club gates cortex-wide communication. Human Brain Mapping, 39(3), 1246–1262. 10.1002/hbm.23913

Shine, J. M., Bissett, P. G., Bell, P. T., Koyejo, O., Balsters, J. H., Gorgolewski, K. J., Moodie, C. A., & Poldrack, R. A. (2016). The Dynamics of Functional Brain Networks: Integrated Network States during Cognitive Task Performance. Neuron, 92(2), 544–554. 10.1016/j.neuron.2016.09.018

Sorrentino, P., Seguin, C., Rucco, R., Liparoti, M., Troisi Lopez, E., Bonavita, S., Quarantelli, M., Sorrentino, G., Jirsa, V., & Zalesky, A. (2021). The structural connectome constrains fast brain dynamics. ELife, 10, e67400. 10.7554/eLife.67400

Sporns, O., & Betzel, R. F. (2016). Modular Brain Networks. Annual Review of Psychology, 67(1), 613–640. 10.1146/annurev-psych-122414-033634

Sporns, O., & Zwi, J. D. (2004). The Small World of the Cerebral Cortex. Neuroinformatics, 2(2), 145–162. 10.1385/NI:2:2:145

Stiso, J., & Bassett, D. S. (2018). Spatial Embedding Imposes Constraints on Neuronal Network Architectures. Trends in Cognitive Sciences, 22(12), 1127–1142. 10.1016/j.tics.2018.09.007

Su, W., Guo, Q., Li, Y., Zhang, K., Zhang, Y., & Chen, Q. (2020). Momentary lapses of attention in multisensory environment. Cortex, 131, 195–209. 10.1016/j.cortex.2020.07.014

Suárez, L. E., Markello, R. D., Betzel, R. F., & Misic, B. (2020). Linking Structure and Function in Macroscale Brain Networks. Trends in Cognitive Sciences, 24(4), 302–315. 10.1016/j.tics.2020.01.008

Suweis, S., Tu, C., Rocha, R. P., Zampieri, S., Zorzi, M., & Corbetta, M. (2019). Brain controllability: Not a slam dunk yet. NeuroImage, 200, 552–555. 10.1016/j.neuroimage.2019.07.012

Towlson, E. K., Vértes, P. E., Ahnert, S. E., Schafer, W. R., & Bullmore, E. T. (2013). The Rich Club of the *C. elegans* Neuronal Connectome. The Journal of Neuroscience, 33(15), 6380–6387. 10.1523/JNEUROSCI.3784-12.2013

Tu, C., Rocha, R. P., Corbetta, M., Zampieri, S., Zorzi, M., & Suweis, S. (2018). Warnings and caveats in brain controllability. NeuroImage, 176, 83–91. 10.1016/j.neuroimage.2018.04.010

Valk, S. L., Xu, T., Paquola, C., Park, B., Bethlehem, R. A. I., Vos de Wael, R., Royer, J., Masouleh, S. K., Bayrak, Ş., Kochunov, P., Yeo, B. T. T., Margulies, D., Smallwood, J., Eickhoff, S. B., & Bernhardt, B. C. (2022). Genetic and phylogenetic uncoupling of structure and function in human transmodal cortex. Nature Communications, 13(1), 2341. 10.1038/s41467-022-29886-1

van den Heuvel, M. P., Kahn, R. S., Goñi, J., & Sporns, O. (2012). High-cost, high-capacity backbone for global brain communication. Proceedings of the National Academy of Sciences, 109(28), 11372–11377. 10.1073/pnas.1203593109

van den Heuvel, M. P., & Sporns, O. (2011). Rich-Club Organization of the Human Connectome. The Journal of Neuroscience, 31(44), 15775–15786. 10.1523/JNEUROSCI.3539-11.2011

van den Heuvel, M. P., & Sporns, O. (2019). A cross-disorder connectome landscape of brain dysconnectivity. Nature Reviews Neuroscience, 20(7), 435–446. 10.1038/s41583-019-0177-6

Van Essen, D. C., Smith, S. M., Barch, D. M., Behrens, T. E. J., Yacoub, E., & Ugurbil, K. (2013). The WU-Minn Human Connectome Project: An overview. NeuroImage, 80, 62–79. 10.1016/j.neuroimage.2013.05.041

Váša, F., Seidlitz, J., Romero-Garcia, R., Whitaker, K. J., Rosenthal, G., Vértes, P. E., Shinn, M., Alexander-Bloch, A., Fonagy, P., Dolan, R. J., Jones, P. B., Goodyer, I. M., the NSPN consortium, Sporns, O., & Bullmore, E. T. (2018). Adolescent Tuning of Association Cortex in Human Structural Brain Networks. Cerebral Cortex, 28(1), 281–294. 10.1093/cercor/bhx249

Wu, L., Caprihan, A., & Calhoun, V. (2021). Tracking spatial dynamics of functional connectivity during a task. NeuroImage, 239, 118310. 10.1016/j.neuroimage.2021.118310

Yeo, B. T., Krienen, F. M., Sepulcre, J., Sabuncu, M. R., Lashkari, D., Hollinshead, M., Roffman, J. L., Smoller, J. W., Zöllei, L., & Polimeni, J. R. (2011). The organization of the human cerebral cortex estimated by intrinsic functional connectivity. Journal of Neurophysiology.

Zink, N., Lenartowicz, A., & Markett, S. (2021). A new era for executive function research: On the transition from centralized to distributed executive functioning. Neuroscience & Biobehavioral Reviews, 124, 235–244. 10.1016/j.neubiorev.2021.02.011

