## Supplemental Files for "Exploring the role of the rich club in network control of neurocognitive states"

#### Supplemental figures

**Supplemental Figure 1**

Results of rank ordering for regional control analyses per task and measure

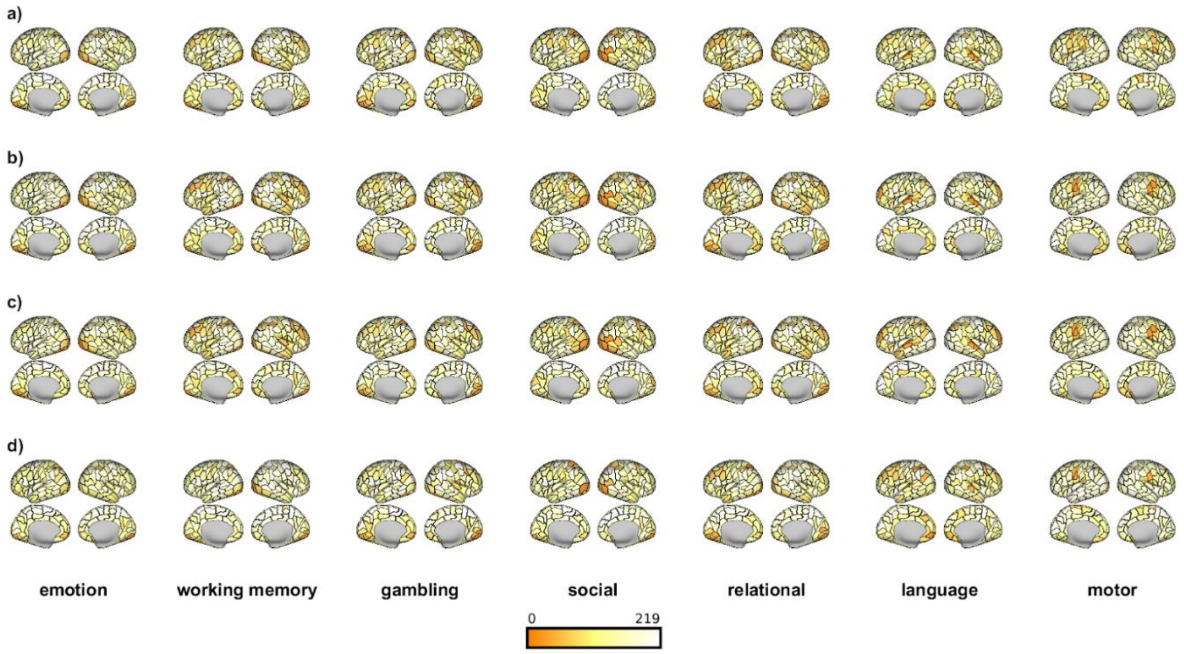


Results of regional control analyses per task and measure. Columns display tasks, rows display measures. Plotted is a region's mean rank in regional control contribution across subjects and tasks, with dark colors indicating low ranks and thus high control contribution, light colors indicating high ranks and thus low control contribution. Black borders indicate different regions as defined in the Lausanne atlas.

a) and b) show mean nodal rank across participants for stability of the stable and instable state, respectively

c) and d) show mean nodal rank across participants for control energy needed to move from the stable to instable and instable to stable state, respectively

**Supplemental Figure 2**

The rich club contributes significantly less to regional control than the periphery in the majority of participants

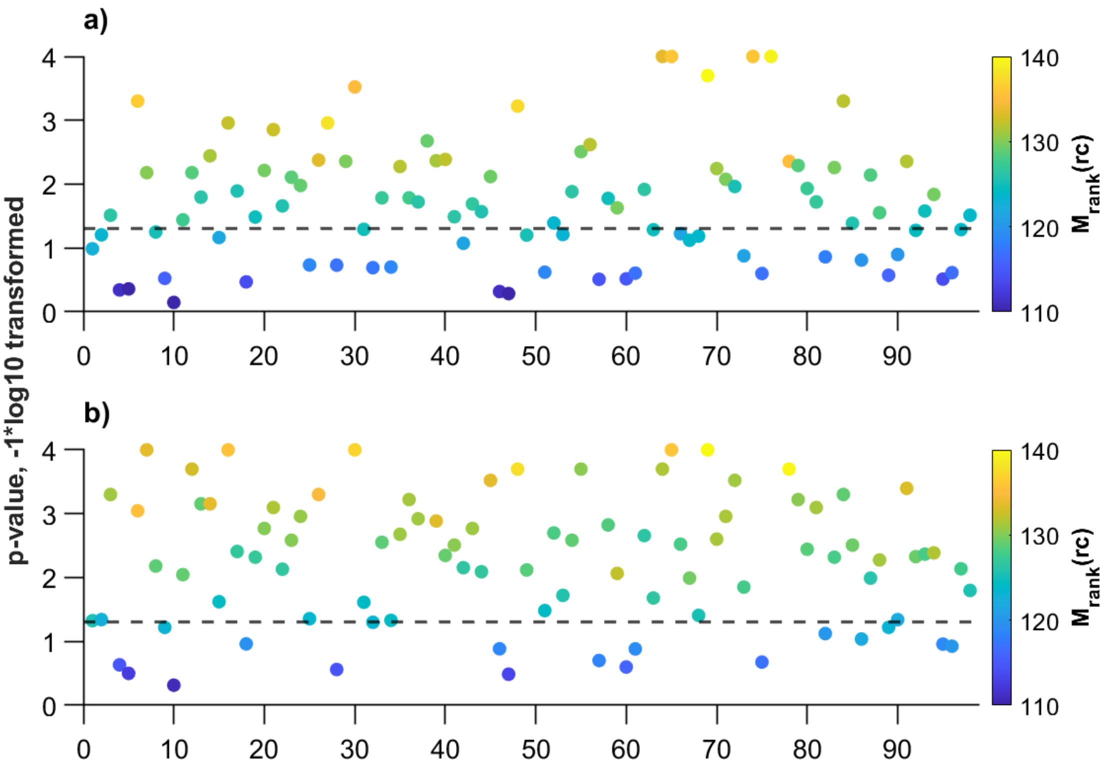


We repeated group-level evaluations of the mean regional control rank of the rich club on a subject level to account for interindividual variation in rich club constitution. The rich club was not significantly involved in control for any participant; for the majority of the sample, it was significantly not-involved. Shown are the log-transformed p-values of comparing the mean regional control rank of subject's rich club to the spin-test based null distribution. Smaller y-values indicate greater p-values. The dashed line indicates a p-value of 0.05; data points above the line show subjects for which the individual rich club was significantly not involved in regional control; data points below the line show subjects for which the individual rich club was not differentially involved in control than a size-matched set of random regions. Colors indicate the mean regional rank, and thus control contribution, of the rich club, with lower rank indicating higher control contributions.

a) Shows values across all tasks and stability measures; the rich club was significantly not involved in regional stability in 85.71% of participants.

b) Shows values across all tasks and energy measures; the rich club was significantly not involved in regional control of state transitions in 73.47% of participants.

**Supplemental Figure 3**

Main effect holds for FA-based and NOS-based connectomes and for a subset of measures in binary connectomes

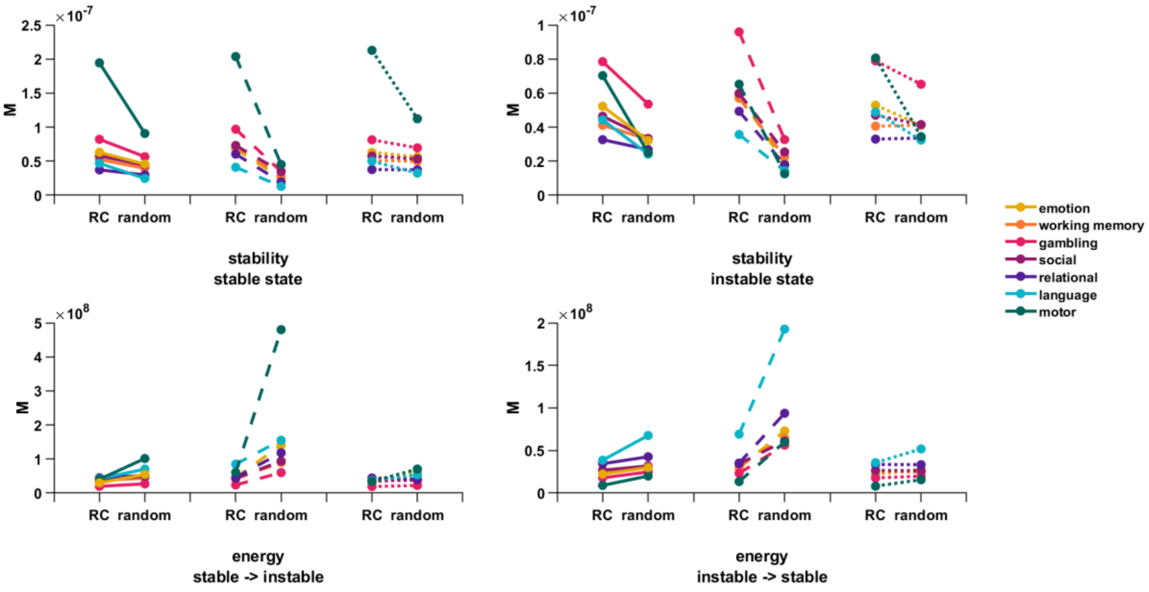


Results of comparing exclusion of rich club (RC) and exclusion of a size-matched set of random regions when connectomes are average FA-based (solid line plots), NOS-based (dashed plots) or binary networks (dotted plots). Displayed are mean NCT values across individuals and for each measure.

*M* = mean value across participants, RC = rich club

**Supplemental Figure 4**

Distributions of NOS-based energy values when including vs. when excluding a major outlier participant

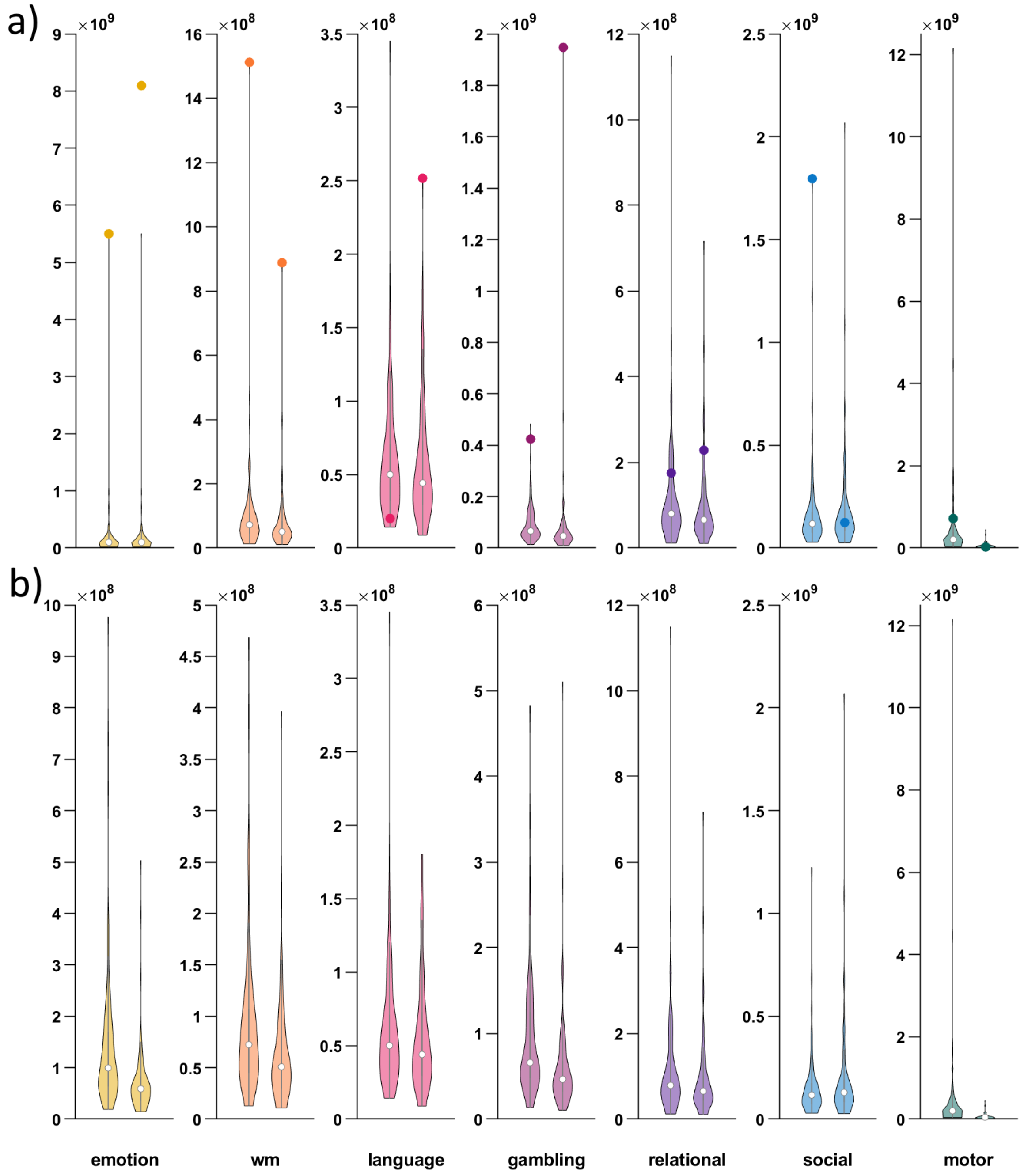


When using a NOS-based connectome for our analyses, we noticed that the control energy values of a single outlier participant distorted results. We here show violins comparing distributions when including the outlier (a) versus when excluding that participant from the sample (b). This outlier participant is excluded for NOS-based results reported in the main text. Shown are distributions of mean energy values for exclusion of randomly selected regions, across participants. For each task, the left plot corresponds to the mean energy values for the transition from the stable to instable state; and the right plot corresponds to the transition from the instable to stable state.

A) Distributions when including the outlier participant; values of the outlier subject are represented with colored dots. This specific individual was qualifying as an outlier in 8 out of 14 possible energy measures.

b) Distributions when the outlier participant shown above was removed from the sample. Statistical evaluations are based on this adjusted sample for NOS-based analyses; we additionally report results for the full sample in Supplemental Tables S11 and S12.

### Supplemental methods

#### HCP acquisition pipeline and processing

Details regarding the acquisition pipeline can be found in van Essen et al. (2013) and Glasser et al. (2013). All MRI data was obtained on a Siemens 3T scanner with a 32 channel head coil.

Regarding diffusion imaging, high-resolution diffusion weighted images (1.25mm isotropic) were obtained using a Stejskal-Tanner (monopolar) diffusion-encoding scheme with a 100 mT/m gradient set and multiband factor of 3, achieving sufficient SNR at 1.25mm resolution with diffusion-weighting up to b=3000 s/mm$^2$. Distortions from eddy currents were addressed by acquiring diffusion data in six series with reversed phase-encoding directions (RL and LR) within each pair. T1- as well as T2-weighted anatomical scans were acquired at resolution of 0.7mm isotropic. The preprocessing pipeline began with intensity normalization of the mean b0 image across the six series. EPI distortions were estimated using FSL's "topup" tool, followed by eddy-current and motion correction with FSL's "eddy" tool. Gradient nonlinearity corrections were applied after running eddy. The corrected diffusion data were then resampled into 1.25mm native structural space and masked. Finally, the corrected b0 image was registered to the T1-weighted structural image using boundary-based registration, transforming the diffusion data to structural space for accurate fiber orientation estimation. Preprocessing analyses relied on FreeSurfer 5.1. (Glasser et al., 2013).

Regarding functional MRI data, grayordinate-based, MSM-All registered results of within-subject task-fMRI analyses were used as provided by the HCP. Data was acquired with a multiband factor of 8, TR = 720ms and a voxel size of 2mm isotropic (van Essen et al., 2013), preprocessed by the consortium itself using a minimal preprocessing pipeline, including gradient unwarping, smoothing to a level of 2mm, motion and distortion correction, registration to MNI152 space and grand-mean intensity normalization (Barch et al., 2013; Glasser et al., 2013). Subsequent within-subject analysis by the HCP included high-pass filtering at 200s and a fixed-effect analysis conducted with the FMRIB Software Library (FSL; Jenkinson et al., 2012) the output is in standard HCP grayordinate space in CIFTI format (Glasser et al., 2013; van Essen et al., 2013, also for further details).

#### Closer description of task paradigms used by the HCP

We used data of all 7 behaviorally constricted tasks acquired by the HCP, namely an emotion matching task, a version of an n-back working memory paradigm, a gambling task for investigation of incentive processing, a language task comparing language to mathematical processing, a motor task eliciting movement, a relational task asking participants to match visual properties of objects, and a social task assessing whether participants thought of objects having a random or social interaction. The 7 tasks provided in the HCP have been chosen with a focus on application to connectivity research and on grounds of their established adequate psychometric criteria, as well as because of their combined ability to investigate a wide array of cognitive functions and activate widespread regions of the brain (Barch et al., 2013). Short descriptions as follows are adapted from information in Barch et al. (2013).

In the emotion task, participants viewed faces displaying different emotions (fearful or neutral) and shapes. Two faces – or objects – were presented at the bottom of the screen, and participants asked to indicate which of the two matches a face or object presented at the top of the screen. Participants were informed through a cue word about which of the two conditions would be following. The task is known to be moderately reliable across time.

The working memory task involved a 2-back and 0-back working memory challenge. Participants were shown blocks of pictures (faces, places, tools, and body parts) and had to indicate if the current image matched one shown two images earlier (2-back) or the currently shown target image (0-back). This task assesses both working memory and category-specific representations. For our investigation, we focused on the working memory aspect of the task, basing brain states category-agnostic on the 0-back and 2-back conditions.

In the gambling task, participants must guess whether a number on a visually presented card is greater or smaller than five to win or lose money. Trials could be rewards, losses, or neutral (with the target number equaling five) and were followed by immediate feedback. The task includes reward blocks (with frequent wins) and loss blocks (with frequent losses). This paradigm is reliably associated with activity in the striatum and other reward-processing brain regions.

The language task contrasts responses when actively listening to a story versus listening to and solving arithmetic problems. The math condition was included as a control since it is unlikely to activate regions involved in semantic processing, and thus constitutes a suitable baseline. In the story condition, participants listened to brief stories and answered forced-choice comprehension questions, with auditory input lasting for 5-9 sentences; in the math condition, participants listened to and solved short arithmetic problems.

To study motion processing, participants performed movements in response to visual cues, including moving their left or right foot, left or right hand, or tongue. Task activity maps on primary motor areas.

The relational processing task required participants to compare pairs of variably shaped and textured objects, judging whether the top and bottom pair match in certain relationships. The task alternated between relational and control conditions, with the relational condition requiring participants to first indicate the dimension of difference between two objects presented at the top, and then judge whether a bottom object pair matches this difference. In the control condition, only one object was presented at the bottom, and participants asked to judge whether this object matched either of the top objects on a previously indicated dimension.

The social cognition paradigm involved watching 20-second videos of objects interacting in ways that imply social interaction (mental condition) versus random movement (random condition). Participants then judged whether the interactions were intentional, not intentional, or whether this was unclear. This task is designed to map onto brain areas involved in mentalization.

#### Supplemental tables

**Table S1**

*Results of repeated measures ANOVA comparing global state stability of the instable and stable state*

| Measures | stability stable state | | stability instable state | | CV[a] | Main effect | |
|---|---|---|---|---|---|---|---|
| | *M* | *SD* | *M* | *SD* | *F*(1,96) | *F*(1,96) | $\eta^2$ |
| Emotion Task | 6.22e-6 | 2.13e-6 | 4.45e-6 | 1.42e-6 | 6.43* | 43.04*** | .31 |
| Working Memory Task | 5.31e-6 | 1.90e-6 | 4.47e-6 | 1.50e-6 | 0.04 | 29.62*** | .24 |
| Language Task | 3.84e-6 | 1.37e-6 | 3.80e-6 | 1.51e-6 | 0.19 | 0.01 | .00 |
| Relational Task | 3.91e-6 | 1.34e-6 | 3.53e-6 | 1.38e-6 | 7.72* | 18.11*** | .16 |
| Social Task | 5.83e-6 | 2.22e-6 | 4.55e-6 | 1.44e-6 | 0.67 | 44.23*** | .32 |
| Gambling Task | 7.76e-6 | 3.15e-6 | 7.24e-6 | 2.84e-6 | 0.98 | 9.75** | .09 |
| Motor Task | 12.83e-6 | 4.71e-6 | 4.25e-6 | 1.85e-6 | 0.00 | 221.59*** | .70 |

*Note.* This table demonstrates means and standard deviations of measures for stability of the stable as well as instable state, as well as corresponding test statistics of repeated measures ANOVA significance testing. Means and standard deviations are rounded to increase comparability for the reader.

[a] CV = covariate, here mean difference of activation between the instable and stable state

*** $p < 0.001$, ** $p < 0.01$, * $p < 0.05$

**Table S2**

*Results of repeated measures ANOVA comparing global control energy transitioning between the stable and instable state*

| Measures | stable -> instable state | | instable -> stable state | | CV | Main effect | |
|---|---|---|---|---|---|---|---|
| | *M* | *SD* | *M* | *SD* | *F*(1,96) | *F*(1,96) | $\eta^2$ |
| Emotion Task | 4.21e+5 | 2.35e+5 | 2.37e+5 | 1.09e+5 | 25.56*** | 11.28** | .11 |
| Working Memory Task | 3.30e+5 | 1.24e+5 | 2.46e+5 | 1.04e+5 | 0.81 | 30.83*** | .24 |
| Language Task | 4.97e+5 | 1.76e+5 | 4.78e+5 | 1.67e+5 | 0.72 | 0.00 | .00 |
| Relational Task | 4.09e+5 | 1.78e+5 | 3.35e+5 | 2.09e+5 | 11.78*** | 26.63*** | .22 |
| Social Task | 3.78e+5 | 1.77e+5 | 2.58e+5 | 1.74e+5 | 1.77 | 50.39*** | .34 |
| Gambling Task | 2.09e+5 | 0.83e+5 | 1.87e+5 | 0.79e+5 | 0.03 | 5.57* | .05 |
| Motor Task | 5.96e+5 | 3.28e+5 | 1.53e+5 | 1.15e+5 | 8.64** | 104.57*** | .52 |

*Note.* This table demonstrates means and standard deviations of measures for control energy for the transition between the stable and instable state, as well as corresponding test statistics of repeated measures ANOVA significance testing. Means and standard deviations are rounded to increase comparability for the reader.

[a] CV = covariate, here mean difference of activation between the instable and stable state

*** $p < 0.001$, ** $p < 0.01$, * $p < 0.05$

**Table S3**

*Results of repeated measures ANOVA comparing state stability between exclusion of rich club members and a size-matched set of semi-randomly selected regions*

| Measures | RC excluded | | Random excluded | | CV | Main effect | |
|---|---|---|---|---|---|---|---|
| | *M* | *SD* | *M* | *SD* | *F*(1,96) | *F*(1,96) | $\eta^2$ |
| Stability stable state | | | | | | | |
| Emotion Task | 6.31e-8 | 3.26e-8 | 4.57e-8 | 1.62e-8 | 0.03 | 15.35*** | .14 |
| Working Memory Task | 5.25e-8 | 3.00e-8 | 3.96e-8 | 1.41e-8 | 0.00 | 12.21*** | .11 |
| Language Task | 4.69e-8 | 2.75e-8 | 2.47e-8 | 0.91e-8 | 0.16 | 32.06*** | .25 |
| Relational Task | 3.73e-8 | 1.77e-8 | 2.99e-8 | 1.06e-8 | 0.13 | 14.86*** | .13 |
| Social Task | 5.78e-8 | 3.07e-8 | 4.43e-8 | 1.82e-8 | 0.05 | 15.18*** | .14 |
| Gambling Task | 8.22e-8 | 4.24e-8 | 5.66e-8 | 2.58e-8 | 0.41 | 16.75*** | .15 |
| Motor Task | 19.44e-8 | 12.28e-8 | 9.08e-8 | 3.75e-8 | 0.40 | 54.57*** | .36 |
| Stability instable state | | | | | | | |
| Emotion Task | 5.23e-8 | 2.67e-8 | 3.22e-8 | 1.10e-8 | 0.18 | 35.29*** | .27 |
| Working Memory Task | 4.11e-8 | 2.08e-8 | 3.29e-8 | 1.15e-8 | 0.00 | 12.40** | .11 |
| Language Task | 4.43e-8 | 2.48e-8 | 2.41e-8 | 0.93e-8 | 0.16 | 35.67*** | .27 |
| Relational Task | 3.26e-8 | 1.87e-8 | 2.68e-8 | 1.06e-8 | 0.05 | 10.29** | .10 |
| Social Task | 4.64e-8 | 2.26e-8 | 3.34e-8 | 1.19e-8 | 0.02 | 23.60*** | .20 |
| Gambling Task | 7.85e-8 | 4.32e-8 | 5.35e-8 | 2.33e-8 | 1.10 | 14.93*** | .13 |
| Motor Task | 7.03e-8 | 5.06e-8 | 2.49e-8 | 1.21e-8 | 7.42** | 83.08*** | .46 |

*Note.* This table demonstrates means and standard deviations of measures for stability of the instable and stable state, as well as corresponding test statistics of repeated measures ANOVA significance testing. Means and standard deviations are rounded to increase comparability for the reader.

[a] CV = covariate, here mean difference of activation between the instable and stable state

*** $p < 0.001$, ** $p < 0.01$, * $p < 0.05$

**Table S4**

*Results of repeated measures ANOVA comparing control energy between exclusion of rich club members and a size-matched set of semi-randomly selected regions*

| Measures | RC excluded | | Random excluded | | CV | Main effect | |
|---|---|---|---|---|---|---|---|
| | *M* | *SD* | *M* | *SD* | *F*(1,96) | *F*(1,96) | $\eta^2$ |
| Energy stable -> instable | | | | | | | |
| Emotion Task | 3.01e+7 | 1.68e+7 | 5.45e+7 | 2.99e+7 | 12.61*** | 31.99*** | .25 |
| Working Memory Task | 3.72e+7 | 1.62e+7 | 4.37e+7 | 1.70e+7 | 1.40 | 4.49* | .04 |
| Language Task | 4.22e+7 | 2.34e+7 | 7.03e+7 | 2.55e+7 | 1.76 | 53.09*** | .36 |
| Relational Task | 4.51e+7 | 2.05e+7 | 5.17e+7 | 2.60e+7 | 0.03 | 5.42* | .05 |
| Social Task | 3.56e+7 | 2.02e+7 | 5.00e+7 | 2.26e+7 | 5.67* | 17.15*** | .15 |
| Gambling Task | 1.98e+7 | 1.07e+7 | 2.71e+7 | 1.08e+7 | 2.23 | 18.93*** | .16 |
| Motor Task | 4.05e+7 | 3.37e+7 | 10.14e+7 | 5.43e+7 | 1.53 | 125.75*** | .57 |
| Energy instable -> stable | | | | | | | |
| Emotion Task | 2.23e+7 | 1.30e+7 | 3.00e+7 | 1.35e+7 | 0.94 | 13.51*** | .12 |
| Working Memory Task | 2.56e+7 | 1.41e+7 | 3.13e+7 | 1.25e+7 | 0.79 | 20.75*** | .18 |
| Language Task | 3.86e+7 | 2.00e+7 | 6.76e+7 | 2.44e+7 | 4.66* | 72.88*** | .43 |
| Relational Task | 3.45e+7 | 2.39e+7 | 4.26e+7 | 2.80e+7 | 1.57 | 4.94* | .05 |
| Social Task | 2.68e+7 | 1.67e+7 | 3.19e+7 | 1.75e+7 | 0.23 | 10.94** | .10 |
| Gambling Task | 1.79e+7 | 0.95e+7 | 2.48e+7 | 1.05e+7 | 4.35* | 18.61*** | .16 |
| Motor Task | 0.90e+7 | 0.77e+7 | 2.00e+7 | 1.11e+7 | 10.45** | 103.08*** | .52 |

*Note.* This table demonstrates means and standard deviations of measures for control energy for the transition between the stable and instable state, as well as corresponding test statistics of repeated measures ANOVA significance testing. Means and standard deviations are rounded to increase comparability for the reader.

[a] CV = covariate, here mean difference of activation between the stable and instable state

*** $p < 0.001$, ** $p < 0.01$, * $p < 0.05$

**Table S5**

*Results of repeated measures ANOVA comparing stability of the stable state between exclusion of rich club members and a size-matched set of semi-randomly selected regions for various settings of the time horizon parameter T*

| From: | RC excluded | | Random excluded | | CV | Main effect | |
|---|---|---|---|---|---|---|---|
| | *M* | *SD* | *M* | *SD* | *F*(1,96) | *F*(1,96) | $\eta^2$ |
| Emotion Task | | | | | | | |
| T = 0.5 | 1.72e-08 | 0.89e-08 | 1.23e-08 | 0.43e-08 | 0.00 | 16.89*** | .15 |
| T = 1.5 | 12.50e-08 | 6.41e-08 | 9.02e-08 | 3.16e-08 | 0.01 | 15.59*** | .14 |
| T = 2 | 1.91e-07 | 0.98e-07 | 1.39e-07 | 0.49e-07 | 0.03 | 15.45*** | .14 |
| T = 2.5 | 2.54e-07 | 1.29e-07 | 1.85e-07 | 0.69e-07 | 0.03 | 14.61*** | .13 |
| T = 3 | 3.12e-07 | 1.59e-07 | 2.29e-07 | 0.85e-07 | 0.06 | 14.40*** | .13 |
| Working Memory Task | | | | | | | |
| T = 0.5 | 1.43e-08 | 0.82e-08 | 1.07e-08 | 0.40e-08 | 0.00 | 13.46*** | .12 |
| T = 1.5 | 10.40e-08 | 5.94e-08 | 7.81e-08 | 2.86e-08 | 0.00 | 13.17*** | .12 |
| T = 2 | 1.59e-07 | 0.91e-07 | 1.20e-07 | 0.44e-07 | 0.00 | 12.73*** | .12 |
| T = 2.5 | 2.11e-07 | 1.21e-07 | 1.61e-07 | 0.58e-07 | 0.00 | 12.16*** | .11 |
| T = 3 | 2.60e-07 | 1.49e-07 | 1.97e-07 | 0.73e-07 | 0.00 | 12.47*** | .12 |
| Language Task | | | | | | | |
| T = 0.5 | 1.28e-08 | 0.75e-08 | 0.67e-08 | 0.26e-08 | 0.15 | 32.71*** | .25 |
| T = 1.5 | 9.28e-08 | 5.43e-08 | 4.90e-08 | 1.81e-08 | 0.13 | 33.63*** | .26 |
| T = 2 | 1.42e-07 | 0.83e-07 | 0.76e-07 | 0.28e-07 | 0.19 | 31.45*** | .25 |
| T = 2.5 | 1.88e-07 | 1.10e-07 | 1.01e-07 | 0.37e-07 | 0.52 | 30.78*** | .24 |
| T = 3 | 2.31e-07 | 1.34e-07 | 1.24e-07 | 0.45e-07 | 0.54 | 30.58*** | .24 |
| Relational Task | | | | | | | |
| T = 0.5 | 1.02e-08 | 0.49e-08 | 0.81e-08 | 0.29e-08 | 0.28 | 16.07*** | .14 |
| T = 1.5 | 7.38e-08 | 3.48e-08 | 5.91e-08 | 2.09e-08 | 0.27 | 15.87*** | .14 |
| T = 2 | 1.13e-07 | 0.53e-07 | 0.90e-07 | 0.31e-07 | 0.26 | 16.95*** | .15 |
| T = 2.5 | 1.50e-07 | 0.70e-07 | 1.20e-07 | 0.42e-07 | 0.26 | 16.77*** | .15 |
| T = 3 | 1.84e-07 | 0.85e-07 | 1.48e-07 | 0.51e-07 | 0.26 | 16.67*** | .15 |
| Social Task | | | | | | | |
| T = 0.5 | 1.58e-08 | 0.84e-08 | 1.19e-08 | 0.49e-08 | 0.03 | 17.75*** | .16 |
| T = 1.5 | 11.44e-08 | 6.05e-08 | 8.70e-08 | 3.59e-08 | 0.05 | 17.42*** | .15 |
| T = 2 | 1.75e-07 | 0.92e-07 | 1.33e-07 | 0.55e-07 | 0.05 | 17.20*** | .15 |
| T = 2.5 | 2.32e-07 | 1.22e-07 | 1.78e-07 | 0.73e-07 | 0.05 | 17.03*** | .15 |
| T = 3 | 2.85e-07 | 1.50e-07 | 2.19e-07 | 0.90e-07 | 0.06 | 16.94*** | .15 |
| Gambling Task | | | | | | | |
| T = 0.5 | 2.24e-08 | 1.17e-08 | 1.54e-08 | 0.73e-08 | 0.55 | 15.01*** | .14 |
| T = 1.5 | 16.24e-08 | 8.32e-08 | 11.27e-08 | 5.11e-08 | 0.35 | 17.27*** | .15 |
| T = 2 | 2.48e-07 | 1.26e-07 | 1.73e-07 | 0.82e-07 | 0.64 | 14.64*** | .13 |
| T = 2.5 | 3.29e-07 | 1.67e-07 | 2.29e-07 | 1.05e-07 | 0.41 | 16.81*** | .15 |
| T = 3 | 4.03e-07 | 2.04e-07 | 2.83e-07 | 1.31e-07 | 0.46 | 16.85*** | .15 |
| Motor Task | | | | | | | |
| T = 0.5 | 5.31e-08 | 3.39e-08 | 2.44e-08 | 0.96e-08 | 0.47 | 53.80*** | .36 |
| T = 1.5 | 38.47e-08 | 24.03e-08 | 17.83e-08 | 7.02e-08 | 0.44 | 55.49*** | .37 |
| T = 2 | 5.88e-07 | 3.64e-07 | 2.74e-07 | 1.08e-07 | 0.42 | 56.22*** | .37 |
| T = 2.5 | 7.82e-07 | 4.81e-07 | 3.65e-07 | 1.44e-07 | 0.41 | 56.73*** | .37 |
| T = 3 | 9.60e-07 | 5.88e-07 | 4.49e-07 | 1.77e-07 | 0.39 | 57.09*** | .37 |

*Note.* This table demonstrates means and standard deviations of measures for stability of the stable state, as well as corresponding test statistics of repeated measures ANOVA significance testing. Means and standard deviations are rounded to increase comparability for the reader.

[a] CV = covariate, here mean difference of activation between the instable and stable state

*** $p < 0.001$, ** $p < 0.01$, * $p < 0.05$

**Table S6**

*Results of repeated measures ANOVA comparing stability of the instable state between exclusion of rich club members and a size-matched set of semi-randomly selected regions for various settings of the time horizon parameter T*

| From: | RC excluded | | Random excluded | | CV | Main effect | |
|---|---|---|---|---|---|---|---|
| | *M* | *SD* | *M* | *SD* | $F(1,96)$ | $F(1,96)$ | $\eta^2$ |
| Emotion Task | | | | | | | |
| T = 0.5 | 1.43e-08 | 0.73e-08 | 0.87e-08 | 0.30e-08 | 0.19 | 35.10*** | .27 |
| T = 1.5 | 10.34e-08 | 5.25e-08 | 6.34e-08 | 2.12e-08 | 0.14 | 34.83*** | .27 |
| T = 2 | 1.58e-07 | 0.80e-07 | 0.98e-07 | 0.34e-07 | 0.12 | 34.21*** | .26 |
| T = 2.5 | 2.10e-07 | 1.06e-07 | 1.29e-07 | 0.45e-07 | 0.17 | 35.51*** | .27 |
| T = 3 | 2.57e-07 | 1.29e-07 | 1.61e-07 | 0.55e-07 | 0.13 | 33.68*** | .26 |
| Working Memory Task | | | | | | | |
| T = 0.5 | 1.12e-08 | 0.57e-08 | 0.89e-08 | 0.31e-08 | 0.01 | 14.00*** | .13 |
| T = 1.5 | 8.13e-08 | 4.09e-08 | 6.49e-08 | 2.30e-08 | 0.01 | 13.46*** | .12 |
| T = 2 | 1.24e-07 | 0.62e-07 | 0.99e-07 | 0.35e-07 | 0.03 | 13.39*** | .12 |
| T = 2.5 | 1.65e-07 | 0.82e-07 | 1.33e-07 | 0.46e-07 | 0.03 | 12.85*** | .12 |
| T = 3 | 2.02e-07 | 1.01e-07 | 1.63e-07 | 0.58e-07 | 0.05 | 13.09*** | .12 |
| Language Task | | | | | | | |
| T = 0.5 | 1.21e-08 | 0.68e-08 | 0.65e-08 | 0.256e-08 | 0.16 | 36.05*** | .27 |
| T = 1.5 | 8.76e-08 | 4.89e-08 | 4.77e-08 | 1.73e-08 | 0.20 | 34.83*** | .27 |
| T = 2 | 1.34e-07 | 0.74e-07 | 0.74e-07 | 0.28e-07 | 0.19 | 34.34*** | .26 |
| T = 2.5 | 1.78e-07 | 0.99e-07 | 0.98e-07 | 0.36e-07 | 0.61 | 30.87*** | .24 |
| T = 3 | 2.19e-07 | 1.21e-07 | 1.21e-07 | 0.45e-07 | 0.64 | 30.64*** | .24 |
| Relational Task | | | | | | | |
| T = 0.5 | 0.89e-08 | 0.51e-08 | 0.73e-08 | 0.29e-08 | 0.16 | 10.79** | .10 |
| T = 1.5 | 6.46e-08 | 3.69e-08 | 5.33e-08 | 2.14e-08 | 0.12 | 10.51** | .10 |
| T = 2 | 0.99e-07 | 0.56e-07 | 0.81e-07 | 0.31e-07 | 0.09 | 11.00** | .10 |
| T = 2.5 | 1.31e-07 | 0.75e-07 | 1.08e-07 | 0.42e-07 | 0.09 | 10.75** | .10 |
| T = 3 | 1.61e-07 | 0.92e-07 | 1.33e-07 | 0.52e-07 | 0.09 | 10.59** | .10 |
| Social Task | | | | | | | |
| T = 0.5 | 1.27e-08 | 0.62e-08 | 0.89e-08 | 0.32e-08 | 0.01 | 27.01*** | .22 |
| T = 1.5 | 9.19e-08 | 4.46e-08 | 6.53e-08 | 2.31e-08 | 0.01 | 26.84*** | .22 |
| T = 2 | 1.40e-07 | 0.68e-07 | 1.00e-07 | 0.35e-07 | 0.01 | 26.70**** | .22 |
| T = 2.5 | 1.87e-07 | 0.90e-07 | 1.33e-07 | 0.47e-07 | 0.01 | 26.61*** | .22 |
| T = 3 | 2.29e-07 | 1.10e-07 | 1.64e-07 | 0.58e-07 | 0.02 | 26.59*** | .22 |
| Gambling Task | | | | | | | |
| T = 0.5 | 2.14e-08 | 1.19e-08 | 1.46e-08 | 0.66e-08 | 1.34 | 14.01*** | .13 |
| T = 1.5 | 15.50e-08 | 8.48e-08 | 10.68e-08 | 4.72e-08 | 1.09 | 14.78*** | .13 |
| T = 2 | 2.36e-07 | 1.28e-07 | 1.64e-07 | 0.74e-07 | 1.37 | 13.53*** | .12 |
| T = 2.5 | 3.14e-07 | 1.70e-07 | 2.16e-07 | 0.94e-07 | 1.10 | 15.24*** | .14 |
| T = 3 | 3.85e-07 | 2.07e-07 | 2.67e-07 | 1.18e-07 | 1.26 | 15.10*** | .14 |

| | | | | | | | |
|---|---|---|---|---|---|---|---|
| Motor Task | | | | | | | |
| T = 0.5 | 1.92e-08 | 1.40e-08 | 0.68e-08 | 0.33e-08 | 7.40** | 83.70*** | .47 |
| T = 1.5 | 13.89e-08 | 9.89e-08 | 4.96e-08 | 2.41e-08 | 7.46** | 86.64*** | .47 |
| T = 2 | 2.12e-07 | 1.50e-07 | 0.76e-07 | 0.37e-07 | 7.47** | 87.93*** | .48 |
| T = 2.5 | 2.82e-07 | 1.98e-07 | 1.02e-07 | 0.50e-07 | 7.46** | 88.88*** | .48 |
| T = 3 | 3.46e-07 | 2.41e-07 | 1.25e-07 | 0.61e-07 | 7.45** | 89.61*** | .48 |

*Note.* This table demonstrates means and standard deviations of measures for stability of the instable state, as well as corresponding test statistics of repeated measures ANOVA significance testing. Means and standard deviations are rounded to increase comparability for the reader.

[a] CV = covariate, here mean difference of activation between the instable and stable state

*** $p < 0.001$, ** $p < 0.01$, * $p < 0.05$

**Table S7**

*Results of repeated measures ANOVA comparing control energy when traversing from the stable to the instable state for various settings of the time horizon parameter T; comparing exclusion of rich club members and a size-matched set of semi-randomly selected regions*

| From: | RC excluded | | Random excluded | | CV | Main effect | |
|---|---|---|---|---|---|---|---|
| | *M* | *SD* | *M* | *SD* | $F(1,96)$ | $F(1,96)$ | $\eta^2$ |
| Emotion Task | | | | | | | |
| T = 0.5 | 17.21e+07 | 11.51e+07 | 37.79e+07 | 22.52e+07 | 22.42*** | 40.31*** | .30 |
| T = 1.5 | 1.34e+07 | 0.70e+07 | 2.24e+07 | 1.16e+07 | 9.31** | 27.38*** | .22 |
| T = 2 | 8.37e+06 | 4.25e+06 | 13.34e+06 | 6.83e+06 | 6.76* | 26.53*** | .22 |
| T = 2.5 | 6.14e+06 | 3.08e+06 | 9.62e+06 | 4.83e+06 | 5.24* | 28.06*** | .23 |
| T = 3 | 4.93e+06 | 2.46e+06 | 7.52e+06 | 3.75e+06 | 5.13* | 24.25*** | .20 |
| Working Memory Task | | | | | | | |
| T = 0.5 | 20.19e+07 | 9.20e+07 | 25.38e+07 | 11.58e+07 | 5.02* | 6.14* | .06 |
| T = 1.5 | 1.68e+07 | 0.74e+07 | 1.95e+07 | 0.70e+07 | 0.27 | 5.86* | .06 |
| T = 2 | 10.49e+06 | 4.68e+06 | 12.04e+06 | 4.22e+06 | 0.06 | 5.51* | .05 |
| T = 2.5 | 7.70e+06 | 3.46e+06 | 8.75e+06 | 3.06e+06 | 0.05 | 4.69* | .05 |
| T = 3 | 6.20e+06 | 2.80e+06 | 7.08e+06 | 2.60e+06 | 0.00 | 5.86* | .06 |
| Language Task | | | | | | | |
| T = 0.5 | 29.67e+07 | 16.77e+07 | 53.02e+07 | 19.42e+07 | 3.99* | 54.90*** | .36 |
| T = 1.5 | 1.76e+07 | 0.99e+07 | 2.86e+07 | 1.05e+07 | 0.86 | 53.43*** | .36 |
| T = 2 | 10.67e+06 | 6.04e+06 | 17.04e+06 | 6.39e+06 | 0.49 | 52.58*** | .35 |
| T = 2.5 | 7.73e+06 | 4.40e+06 | 12.24e+06 | 4.64e+06 | 2.99 | 37.21*** | .28 |
| T = 3 | 6.18e+06 | 3.52e+06 | 9.73e+06 | 3.70e+06 | 2.91 | 36.54*** | .28 |
| Relational Task | | | | | | | |
| T = 0.5 | 22.84e+07 | 11.51e+07 | 27.25e+07 | 13.68e+07 | 1.43 | 4.53* | .05 |
| T = 1.5 | 2.09e+07 | 0.94e+07 | 2.36e+07 | 1.16e+07 | 0.55 | 6.11* | .06 |
| T = 2 | 13.19e+06 | 5.93e+06 | 14.77e+06 | 6.45e+06 | 0.70 | 6.79* | .07 |
| T = 2.5 | 9.75e+06 | 4.40e+06 | 10.88e+06 | 4.77e+06 | 0.84 | 6.68* | .07 |
| T = 3 | 7.87e+06 | 3.56e+06 | 8.77e+06 | 3.86e+06 | 0.91 | 6.60* | .06 |
| Social Task | | | | | | | |
| T = 0.5 | 22.20e+07 | 14.73e+07 | 34.19e+07 | 19.66e+07 | 15.49*** | 18.55*** | .16 |
| T = 1.5 | 1.54e+07 | 0.84e+07 | 2.11e+07 | 0.89e+07 | 2.28 | 18.97*** | .17 |
| T = 2 | 9.47e+06 | 5.11e+06 | 1.27e+07 | 0.52e+07 | 1.32 | 18.02*** | .16 |

| | | | | | | | |
|---|---|---|---|---|---|---|---|
| T = 2.5 | 6.91e+06 | 3.72e+06 | 9.14e+06 | 3.71e+06 | 0.92 | 17.27*** | .15 |
| T = 3 | 5.54e+06 | 2.97e+06 | 7.27e+06 | 2.94e+06 | 0.73 | 16.79*** | .15 |
| Gambling Task | | | | | | | |
| T = 0.5 | 10.86e+07 | 6.95e+07 | 16.18e+07 | 7.98e+07 | 9.08** | 14.52*** | .13 |
| T = 1.5 | 0.90e+07 | 0.47e+07 | 1.21e+07 | 0.47e+07 | 0.39 | 24.01*** | .20 |
| T = 2 | 5.69e+06 | 2.90e+06 | 7.50e+06 | 2.80e+06 | 0.22 | 24.85*** | .21 |
| T = 2.5 | 4.20e+06 | 2.13e+06 | 5.52e+06 | 2.07e+06 | 0.34 | 24.57*** | .20 |
| T = 3 | 3.39e+06 | 1.71e+06 | 4.44e+06 | 1.68e+06 | 0.22 | 26.13*** | .21 |
| Motor Task | | | | | | | |
| T = 0.5 | 32.23e+07 | 29.49e+07 | 85.16e+07 | 47.23e+07 | 0.43 | 120.67*** | .56 |
| T = 1.5 | 1.52e+07 | 1.19e+07 | 3.66e+07 | 1.88e+07 | 1.50 | 121.32*** | .56 |
| T = 2 | 8.60e+06 | 6.55e+06 | 20.26e+06 | 10.26e+06 | 1.69 | 121.00*** | .56 |
| T = 2.5 | 5.99e+06 | 4.47e+06 | 13.91e+06 | 6.97e+06 | 1.77 | 120.57*** | .56 |
| T = 3 | 4.67e+06 | 3.44e+06 | 10.75e+06 | 5.36e+06 | 1.80 | 120.18*** | .56 |

*Note.* This table demonstrates means and standard deviations of measures for energy needed for transitions from the stable to the instable state, as well as corresponding test statistics of repeated measures ANOVA significance testing. Means and standard deviations are rounded to increase comparability for the reader.

[a] CV = covariate, here mean difference of activation between the instable and stable state

*** $p < 0.001$, ** $p < 0.01$, * $p < 0.05$

**Table S8**

*Results of repeated measures ANOVA comparing control energy when traversing from the instable to the stable state for various settings of the time horizon parameter T; comparing exclusion of rich club members and a size-matched set of semi-randomly selected regions*

| From: | RC excluded | | Random excluded | | CV | Main effect | |
|---|---|---|---|---|---|---|---|
| | *M* | *SD* | *M* | *SD* | $F(1,96)$ | $F(1,96)$ | $\eta^2$ |
| Emotion Task | | | | | | | |
| T = 0.5 | 11.91e+07 | 7.90e+07 | 20.82e+07 | 9.97e+07 | 7.04** | 30.47*** | .24 |
| T = 1.5 | 1.05e+07 | 0.58e+07 | 1.34e+07 | 0.59e+07 | 0.23 | 13.39*** | .12 |
| T = 2 | 6.75+06 | 3.62e+06 | 8.45e+06 | 3.77e+06 | 0.26 | 12.27*** | .11 |
| T = 2.5 | 5.03e+06 | 2.66e+06 | 6.33e+06 | 2.86e+06 | 0.17 | 12.54*** | .12 |
| T = 3 | 4.08e+06 | 2.14e+06 | 5.11e+06 | 2.27e+06 | 0.45 | 11.53*** | .11 |
| Working Memory Task | | | | | | | |
| T = 0.5 | 12.14e+07 | 7.29e+07 | 16.67e+07 | 7.81e+07 | 0.00 | 36.06*** | .27 |
| T = 1.5 | 1.25e+07 | 0.67e+07 | 1.49e+07 | 0.56e+07 | 1.67 | 19.43*** | .17 |
| T = 2 | 8.16e+06 | 4.30e+06 | 9.58e+06 | 3.58e+06 | 1.50 | 15.14*** | .14 |
| T = 2.5 | 6.13e+06 | 3.21e+06 | 7.07e+06 | 2.54e+06 | 1.17 | 11.73*** | .11 |
| T = 3 | 5.00e+06 | 2.60e+06 | 5.83e+06 | 2.25e+06 | 1.32 | 13.49*** | .12 |
| Language Task | | | | | | | |
| T = 0.5 | 27.26e+07 | 11.57e+07 | 50.45e+07 | 17.05e+07 | 9.25** | 76.57*** | .44 |
| T = 1.5 | 1.62e+07 | 0.91e+07 | 2.76e+07 | 1.05e+07 | 1.78 | 69.51*** | .42 |
| T = 2 | 9.91e+06 | 5.74e+06 | 16.39e+06 | 6.05e+06 | 2.08 | 58.61*** | .38 |
| T = 2.5 | 7.21e+06 | 4.21e+06 | 11.87e+06 | 4.53e+06 | 4.24* | 44.26*** | .32 |
| T = 3 | 5.77e+06 | 3.39e+06 | 9.45e+06 | 3.62e+06 | 4.19* | 43.51*** | .31 |
| Relational Task | | | | | | | |
| T = 0.5 | 15.54e+07 | 13.79e+07 | 21.06e+07 | 16.98e+07 | 14.64*** | 3.14 | .03 |

| | | | | | | | |
|---|---|---|---|---|---|---|---|
| T = 1.5 | 1.70e+07 | 1.07e+07 | 2.04e+07 | 1.22e+07 | 0.06 | 6.37* | .06 |
| T = 2 | 11.04e+06 | 6.67e+06 | 13.10e+06 | 7.05e+06 | 0.00 | 7.87** | .08 |
| T = 2.5 | 8.30e+06 | 4.91e+06 | 9.76e+06 | 5.15e+06 | 0.08 | 7.97** | .08 |
| T = 3 | 6.76e+06 | 3.96e+06 | 7.91e+06 | 4.13e+06 | 0.16 | 8.04** | .08 |
| Social Task | | | | | | | |
| T = 0.5 | 16.22e+07 | 13.41e+07 | 21.32e+07 | 17.29e+07 | 8.76** | 7.20** | .07 |
| T = 1.5 | 1.21e+07 | 0.68e+07 | 1.44e+07 | 0.68e+07 | 0.29 | 13.71*** | .13 |
| T = 2 | 7.67e+06 | 4.16e+06 | 9.05e+06 | 4.01e+06 | 0.72 | 14.77*** | .13 |
| T = 2.5 | 5.68e+06 | 3.03e+06 | 6.70e+06 | 2.89e+06 | 0.97 | 15.30*** | .14 |
| T = 3 | 4.60e+06 | 2.43e+06 | 5.42e+06 | 2.31e+06 | 1.10 | 15.66*** | .14 |
| Gambling Task | | | | | | | |
| T = 0.5 | 9.55e+07 | 5.75e+07 | 14.50e+07 | 7.84e+07 | 16.44*** | 10.41** | .10 |
| T = 1.5 | 0.83e+07 | 0.43e+07 | 1.12e+07 | 0.45e+07 | 1.56 | 22.37*** | .19 |
| T = 2 | 5.30e+06 | 2.73e+06 | 7.05e+06 | 2.77e+06 | 1.35 | 21.47*** | .18 |
| T = 2.5 | 3.94e+06 | 2.02e+06 | 5.22e+06 | 2.05e+06 | 0.88 | 22.13*** | .19 |
| T = 3 | 3.19e+06 | 1.63e+06 | 4.20e+06 | 1.65e+06 | 0.64 | 23.25*** | .20 |
| Motor Task | | | | | | | |
| T = 0.5 | 10.58e+07 | 11.41e+07 | 28.80e+07 | 17.37e+07 | 5.09* | 127.60*** | .57 |
| T = 1.5 | 0.36e+07 | 0.24e+07 | 0.70e+07 | 0.36e+07 | 5.66* | 96.18*** | .50 |
| T = 2 | 2.23e+06 | 1.35e+06 | 4.23e+06 | 2.07e+06 | 1.84 | 95.95*** | .50 |
| T = 2.5 | 1.67e+06 | 0.97e+06 | 3.15e+06 | 1.49e+06 | 0.64 | 99.94*** | .51 |
| T = 3 | 1.37e+06 | 0.77e+06 | 2.57e+06 | 1.20e+06 | 0.25 | 103.43*** | .52 |

*Note.* This table demonstrates means and standard deviations of measures for energy needed for transitions from the instable to the stable state, as well as corresponding test statistics of repeated measures ANOVA significance testing. Means and standard deviations are rounded to increase comparability for the reader.

[a] CV = covariate, here mean difference of activation between the instable and stable state

*** $p < 0.001$, ** $p < 0.01$, * $p < 0.05$

**Table S9**

*Results of repeated measures ANOVA comparing state stability between exclusion of rich club members and a size-matched set of semi-randomly selected regions; analyses based on a NOS-weighted connectome; outlier not included*

| Measures | RC excluded | | Random excluded | | CV | Main effect | |
|---|---|---|---|---|---|---|---|
| | *M* | *SD* | *M* | *SD* | *F*(1,96) | *F*(1,96) | $\eta^2$ |
| Stability stable state | | | | | | | |
| Emotion Task | 6.82e-08 | 4.99e-08 | 2.51e-08 | 1.65e-08 | 0.06 | 41.25*** | .30 |
| Working Memory Task | 7.20e-08 | 6.10e-08 | 2.77e-08 | 1.87e-08 | 2.16 | 44.35*** | .32 |
| Language Task | 4.11e-08 | 4.17e-08 | 1.31e-08 | 0.85e-08 | 0.01 | 20.36*** | .18 |
| Relational Task | 6.04e-08 | 4.39e-08 | 1.95e-08 | 1.32e-08 | 0.17 | 51.66*** | .35 |
| Social Task | 7.31e-08 | 5.54e-08 | 3.47e-08 | 2.48e-08 | 0.59 | 48.69*** | .34 |
| Gambling Task | 9.70e-08 | 6.59e-08 | 3.49e-08 | 2.42e-08 | 0.03 | 48.28*** | .34 |
| Motor Task | 20.37e-08 | 19.05e-08 | 4.52e-08 | 3.18e-08 | 0.44 | 43.08*** | .31 |
| Stability instable state | | | | | | | |
| Emotion Task | 5.88e-08 | 4.78e-08 | 1.87e-08 | 1.36e-08 | 0.10 | 37.66*** | .28 |
| Working Memory Task | 5.69e-08 | 4.81e-08 | 2.26e-08 | 1.54e-08 | 0.12 | 32.97*** | .26 |
| Language Task | 3.57e-08 | 3.16e-08 | 1.43e-08 | 0.93e-08 | 0.04 | 22.64*** | .19 |
| Relational Task | 4.93e-08 | 3.64e-08 | 1.79e-08 | 1.36e-08 | 0.25 | 52.12*** | .35 |
| Social Task | 5.98e-08 | 4.45e-08 | 2.54e-08 | 1.75e-08 | 0.67 | 46.18*** | .33 |
| Gambling Task | 9.59e-08 | 7.00e-08 | 3.27e-08 | 2.00e-08 | 0.54 | 35.71*** | .27 |
| Motor Task | 6.52e-08 | 5.21e-08 | 1.26e-08 | 1.09e-08 | 2.30 | 72.99*** | .43 |

*Note.* This table demonstrates means and standard deviations of measures for stability of the stable and instable state, as well as corresponding test statistics of repeated measures ANOVA significance testing. Means and standard deviations are rounded to increase comparability for the reader. **Note that one subject was identified as an outlier (all energy values > 2IQR distance from the group) and subsequently excluded from inferential analyses. See Supplemental Figure S4 and Supplemental Tables S11 and S12 for a visualization of the outlier.**

[a] CV = covariate, here mean difference of activation between the stable and instable state

*** $p < 0.001$, ** $p < 0.01$, * $p < 0.05$

**Table S10**

*Results of repeated measures ANOVA comparing control energy between exclusion of rich club members and a size-matched set of semi-randomly selected regions; analyses based on a NOS-weighted connectome; outlier not included*

| Measures | RC excluded | | Random excluded | | CV | Main effect | |
|---|---|---|---|---|---|---|---|
| | *M* | *SD* | *M* | *SD* | *F*(1,96) | *F*(1,96) | $\eta^2$ |
| Energy stable -> instable | | | | | | | |
| Emotion Task | 4.53e+07 | 4.51e+07 | 14.10e+07 | 14.27e+07 | 22.77*** | 4.72* | .05 |
| Working Memory Task | 4.17e+07 | 3.80e+07 | 8.98e+07 | 7.25e+07 | 0.24 | 19.39*** | .17 |
| Language Task | 8.49e+07 | 9.24e+07 | 15.53e+07 | 15.30e+07 | 1.03 | 4.26* | .04 |
| Relational Task | 4.56e+07 | 4.12e+07 | 11.84e+07 | 13.41e+07 | 0.25 | 11.97*** | .11 |
| Social Task | 4.33e+07 | 3.92e+07 | 9.38e+07 | 7.33e+07 | 4.07* | 18.46*** | .16 |
| Gambling Task | 2.39e+07 | 2.13e+07 | 6.00e+07 | 4.45e+07 | 0.17 | 26.17*** | .22 |
| Motor Task | 6.10e+07 | 6.35e+07 | 47.98e+07 | 131.16e+07 | 0.53 | 7.39** | .07 |
| Energy instable -> stable | | | | | | | |
| Emotion Task | 2.94e+07 | 3.12e+07 | 7.2978e+07 | 6.85e+07 | 0.85 | 14.12*** | .13 |
| Working Memory Task | 3.21e+07 | 4.51e+07 | 6.45e+07 | 5.33e+07 | 2.46 | 27.38*** | .22 |
| Language Task | 6.91e+07 | 8.72e+07 | 19.26e+07 | 24.82e+07 | 0.18 | 7.33** | .07 |
| Relational Task | 3.52e+07 | 4.59e+07 | 9.37e+07 | 9.59e+07 | 22.32*** | 7.955** | .08 |
| Social Task | 3.45e+07 | 4.11e+07 | 6.10e+07 | 6.21e+07 | 12.60*** | 5.14* | .05 |
| Gambling Task | 2.33e+07 | 2.30e+07 | 5.63e+07 | 3.80e+07 | 0.04 | 37.23*** | .28 |
| Motor Task | 1.35e+07 | 1.71e+07 | 6.00e+07 | 6.61e+07 | 1.23 | 16.95*** | .15 |

*Note.* This table demonstrates means and standard deviations of measures for control energy for the transition between the stable and instable state, as well as corresponding test statistics of repeated measures ANOVA significance testing. Means and standard deviations are rounded to increase comparability for the reader. **Note that one subject was identified as an outlier (all energy values > 2IQR distance from the group) and subsequently excluded from inferential analyses. See figure Supplemental Figure S4 and Supplemental Tables S11 and S12 for a visualization of the outlier.**

[a] CV = covariate, here mean difference of activation between the stable and instable state.

*** $p < 0.001$, ** $p < 0.01$, * $p < 0.05$

**Table S11**

*Results of repeated measures ANOVA comparing state stability between exclusion of rich club members and a size-matched set of semi-randomly selected regions; analyses based on a NOS-weighted connectome; all 98 subjects included*

| Measures | RC excluded | | Random excluded | | CV | Main effect | |
|---|---|---|---|---|---|---|---|
| | *M* | *SD* | *M* | *SD* | *F*(1,96) | *F*(1,96) | $\eta^2$ |
| Stability stable state | | | | | | | |
| Emotion Task | 6.78e-08 | 4.98e-08 | 2.49e-08 | 1.66e-08 | 0.06 | 41.72*** | .30 |
| Working Memory Task | 7.16e-08 | 6.09e-08 | 2.75e-08 | 1.88e-08 | 2.30 | 44.85*** | .32 |
| Language Task | 4.10e-08 | 4.15e-08 | 1.31e-08 | 0.85e-08 | 0.01 | 20.72*** | .18 |
| Relational Task | 6.00e-08 | 4.38e-08 | 1.93e-08 | 1.32e-08 | 0.14 | 51.91*** | .35 |
| Social Task | 7.29e-08 | 5.52e-08 | 3.44e-08 | 2.49e-08 | 0.61 | 49.97*** | .34 |
| Gambling Task | 9.75e-08 | 6.58e-08 | 3.46e-08 | 2.43e-08 | 0.00 | 51.32*** | .35 |
| Motor Task | 20.40e-08 | 18.96e-08 | 4.49e-08 | 3.17e-08 | 0.48 | 44.75*** | .32 |
| Stability instable state | | | | | | | |
| Emotion Task | 5.86e-08 | 4.76e-08 | 1.85e-08 | 1.36e-08 | 0.10 | 38.28*** | .29 |
| Working Memory Task | 5.66e-08 | 4.79e-08 | 2.24e-08 | 1.54e-08 | 0.14 | 33.35*** | .26 |
| Language Task | 3.63e-08 | 3.20e-08 | 1.42e-08 | 0.93e-08 | 0.07 | 19.89*** | .17 |
| Relational Task | 4.90e-08 | 3.63e-08 | 1.78e-08 | 1.36e-08 | 0.22 | 52.42*** | .35 |
| Social Task | 5.96e-08 | 4.43e-08 | 2.54e-08 | 1.74e-08 | 0.64 | 45.95*** | .32 |
| Gambling Task | 9.60e-08 | 6.96e-08 | 3.26e-08 | 1.99e-08 | 0.50 | 37.32*** | .38 |
| Motor Task | 6.58e-08 | 5.21e-08 | 1.25e-08 | 1.09e-08 | 2.62 | 76.62*** | .44 |

*Note.* This table demonstrates means and standard deviations of measures for stability of the stable and instable state, as well as corresponding test statistics of repeated measures ANOVA significance testing. Means and standard deviations are rounded to increase comparability for the reader.

[a] CV = covariate, here mean difference of activation between the stable and instable state

*** $p < 0.001$, ** $p < 0.01$, * $p < 0.05$

**Table S12**

*Results of repeated measures ANOVA comparing control energy between exclusion of rich club members and a size-matched set of semi-randomly selected regions; analyses based on a NOS-weighted connectome; all 98 subjects included*

| Measures | RC excluded | | Random excluded | | CV | Main effect | |
|---|---|---|---|---|---|---|---|
| | *M* | *SD* | *M* | *SD* | *F*(1,96) | *F*(1,96) | $\eta^2$ |
| Energy stable -> instable | | | | | | | |
| Emotion Task | 4.52e+07 | 4.48e+07 | 19.57e+07 | 55.98e+07 | 0.49 | 1.58 | .02 |
| Working Memory Task | 4.18e+07 | 3.79e+07 | 10.43e+07 | 16.07e+07 | 1.39 | 2.93 | .03 |
| Language Task | 8.41e+07 | 9.22e+07 | 17.20e+07 | 22.50e+07 | 4.32* | 0.61 | .01 |
| Relational Task | 4.56e+07 | 4.10e+07 | 11.90e+07 | 13.36e+07 | 0.22 | 12.60*** | .12 |
| Social Task | 4.33e+07 | 3.90e+07 | 9.72e+07 | 8.02e+07 | 2.66 | 18.35*** | .16 |
| Gambling Task | 2.37e+07 | 2.12e+07 | 5.96e+07 | 4.45e+07 | 0.23 | 25.91*** | .21 |
| Motor Task | 6.05e+07 | 6.34e+07 | 48.22e+07 | 130.51e+07 | 0.56 | 7.71** | .07 |
| Energy instable -> stable | | | | | | | |
| Emotion Task | 2.94e+07 | 3.10e+07 | 15.48e+07 | 81.31e+07 | 0.03 | 1.30 | .01 |
| Working Memory Task | 3.22e+07 | 4.49e+07 | 7.29e+07 | 9.86e+07 | 0.01 | 7.00** | .07 |
| Language Task | 6.89e+07 | 8.68e+07 | 19.19e+07 | 24.70e+07 | 0.15 | 7.58** | .07 |
| Relational Task | 3.52e+07 | 4.57e+07 | 9.51e+07 | 9.63e+07 | 19.96*** | 9.30** | .09 |
| Social Task | 3.43e+07 | 4.09e+07 | 8.02e+07 | 20.03e+07 | 0.16 | 1.99 | .02 |
| Gambling Task | 2.31e+07 | 2.29e+07 | 5.83e+07 | 4.26e+07 | 0.09 | 34.78*** | .27 |
| Motor Task | 1.34e+07 | 1.70e+07 | 5.96e+07 | 6.58e+07 | 1.30 | 17.09*** | .15 |

*Note.* This table demonstrates means and standard deviations of measures for control energy for the transition between the stable and instable state, as well as corresponding test statistics of repeated measures ANOVA significance testing. Means and standard deviations are rounded to increase comparability for the reader.

[a] CV = covariate, here mean difference of activation between the stable and instable state.

*** $p < 0.001$, ** $p < 0.01$, * $p < 0.05$

**Table S13**

*Results of repeated measures ANOVA comparing state stability between exclusion of rich club members and a size-matched set of semi-randomly selected regions; analyses based on a binary connectome*

| Measures | RC excluded | | Random excluded | | CV | Main effect | |
|---|---|---|---|---|---|---|---|
| | *M* | *SD* | *M* | *SD* | *F*(1,96) | *F*(1,96) | $\eta^2$ |
| Stability stable state | | | | | | | |
| Emotion Task | 6.30e-08 | 3.28e-08 | 5.64e-08 | 1.99e-08 | 0.22 | 1.30 | .01 |
| Working Memory Task | 5.11e-08 | 2.85e-08 | 5.01e-08 | 1.81e-08 | 0.01 | 0.14 | .00 |
| Language Task | 5.00e-08 | 2.80e-08 | 3.26e-08 | 1.17e-08 | 1.24 | 16.51*** | .15 |
| Relational Task | 3.77e-08 | 1.78e-08 | 3.74e-08 | 1.31e-08 | 0.07 | 0.14 | .00 |
| Social Task | 5.76e-08 | 3.11e-08 | 5.33e-08 | 2.20e-08 | 0.00 | 1.99 | .02 |
| Gambling Task | 8.15e-08 | 4.28e-08 | 6.96e-08 | 2.92e-08 | 0.01 | 4.12* | .04 |
| Motor Task | 21.28e-08 | 5.57e-08 | 11.23e-08 | 4.44e-08 | 0.10 | 38.50*** | .29 |
| Stability instable state | | | | | | | |
| Emotion Task | 5.28e-08 | 2.70e-08 | 4.12e-08 | 1.39e-08 | 0.01 | 10.60** | .10 |
| Working Memory Task | 4.06e-08 | 2.06e-08 | 4.12e-08 | 1.49e-08 | 0.01 | 0.03 | .00 |
| Language Task | 4.88e-08 | 2.67e-08 | 3.24e-08 | 1.20e-08 | 0.87 | 16.48*** | .15 |
| Relational Task | 3.30e-08 | 1.96e-08 | 3.36e-08 | 1.30e-08 | 0.07 | 0.03 | .00 |
| Social Task | 4.70e-08 | 2.32e-08 | 4.15e-08 | 1.43e-08 | 0.06 | 5.27* | .05 |
| Gambling Task | 7.89e-08 | 4.38e-08 | 6.52e-08 | 2.64e-08 | 0.34 | 4.67* | .05 |
| Motor Task | 8.06e-08 | 14.13e-08 | 3.44e-08 | 1.60e-08 | 6.45* | 76.70*** | .44 |

*Note.* This table demonstrates means and standard deviations of measures for stability of the stable and instable state, as well as corresponding test statistics of repeated measures ANOVA significance testing. Means and standard deviations are shown with four decimal numbers to increase comparability for the reader. Means and standard deviations are rounded to increase comparability for the reader.

[a] CV = covariate, here mean difference of activation between the stable and instable state

*** $p < 0.001$, ** $p < 0.01$, * $p < 0.05$

**Table S14**

*Results of repeated measures ANOVA comparing control energy between exclusion of rich club members and a size-matched set of semi-randomly selected regions; analyses based on a binary connectome*

| Measures | RC excluded | | Random excluded | | CV | Main effect | |
|---|---|---|---|---|---|---|---|
| | *M* | *SD* | *M* | *SD* | *F*(1,96) | *F*(1,96) | $\eta^2$ |
| Energy stable -> instable | | | | | | | |
| Emotion Task | 2.98e+07 | 1.59e+07 | 6.28e+07 | 21.16e+07 | 0.47 | 2.37 | .02 |
| Working Memory Task | 3.71e+07 | 1.62e+07 | 3.72e+07 | 2.95e+07 | 0.18 | 0.09 | .00 |
| Language Task | 3.71e+07 | 2.01e+07 | 5.35e+07 | 2.01e+07 | 1.75 | 18.22*** | .16 |
| Relational Task | 4.43e+07 | 1.94e+07 | 4.03e+07 | 1.69e+07 | 0.40 | 2.17 | .02 |
| Social Task | 3.48e+07 | 2.06e+07 | 3.92e+07 | 1.68e+07 | 1.06 | 1.73 | .02 |
| Gambling Task | 1.91e+07 | 0.97e+07 | 2.24e+07 | 1.22e+07 | 0.46 | 1.81 | .02 |
| Motor Task | 3.40e+07 | 2.76e+07 | 7.05e+07 | 3.47e+07 | 3.84 | 143.45*** | .60 |
| Energy instable -> stable | | | | | | | |
| Emotion Task | 2.19e+07 | 1.29e+07 | 2.80e+07 | 3.37e+07 | 0.24 | 2.40 | .02 |
| Working Memory Task | 2.56e+07 | 1.30e+07 | 2.45e+07 | 1.00e+07 | 0.06 | 0.35 | .00 |
| Language Task | 3.57e+07 | 1.69e+07 | 5.19e+07 | 1.73e+07 | 6.65* | 23.79*** | .20 |
| Relational Task | 3.37e+07 | 2.29e+07 | 3.33e+07 | 1.88e+07 | 0.48 | 0.11 | .00 |
| Social Task | 2.66e+07 | 1.65e+07 | 2.63e+07 | 1.35e+07 | 7.40** | 2.77 | .03 |
| Gambling Task | 1.78e+07 | 0.94e+07 | 1.98e+07 | 0.92e+07 | 0.38 | 1.47 | .02 |
| Motor Task | 0.81e+07 | 0.64e+07 | 1.57e+07 | 0.84e+07 | 9.76** | 118.44*** | .55 |

*Note.* This table demonstrates means and standard deviations of measures for control energy for the transition between the stable and instable state, as well as corresponding test statistics of repeated measures ANOVA significance testing. Means and standard deviations are shown with four decimal numbers to increase comparability for the reader.

[a] CV = covariate, here mean difference of activation between the stable and instable state

*** $p < 0.001$, ** $p < 0.01$, * $p < 0.05$

**Table S15**

*Results of repeated measures ANOVA comparing state stability between exclusion of individual level rich club members and a size-matched set of semi-randomly selected regions*

| Measures | RC excluded | | Random excluded | | CV | Main effect | |
|---|---|---|---|---|---|---|---|
| | *M* | *SD* | *M* | *SD* | *F*(1,96) | *F*(1,96) | $\eta^2$ |
| Stability stable state | | | | | | | |
| Emotion Task | 8.83e-08 | 5.89e-08 | 4.55e-08 | 1.66e-08 | 0.24 | 23.14*** | .19 |
| Working Memory Task | 8.26e-08 | 7.16e-08 | 3.94e-08 | 1.46e-08 | 0.15 | 20.62*** | .18 |
| Language Task | 5.59e-08 | 2.70e-08 | 2.47e-08 | 0.89e-08 | 6.07* | 118.11*** | .55 |
| Relational Task | 5.53e-08 | 3.80e-08 | 3.00e-08 | 1.10e-08 | 1.99 | 39.60*** | .29 |
| Social Task | 7.42e-08 | 4.21e-08 | 4.41e-08 | 1.83e-08 | 0.45 | 45.04*** | .32 |
| Gambling Task | 11.28e-08 | 9.95e-08 | 5.67e-08 | 2.62e-08 | 3.10 | 30.51*** | .24 |
| Motor Task | 22.42e-08 | 12.86e-08 | 9.02e-08 | 3.64e-08 | 0.40 | 78.86*** | .45 |
| Stability instable state | | | | | | | |
| Emotion Task | 6.78e-08 | 3.65e-08 | 3.20e-08 | 1.10e-08 | 0.31 | 43.46*** | .31 |
| Working Memory Task | 5.88e-08 | 3.87e-08 | 3.26e-08 | 1.17e-08 | 0.51 | 31.42*** | .25 |
| Language Task | 5.91e-08 | 3.29e-08 | 2.40e-08 | 0.89e-08 | 1.36 | 76.05*** | .44 |
| Relational Task | 4.71e-08 | 2.82e-08 | 2.70e-08 | 1.12e-08 | 1.84 | 42.10*** | .30 |
| Social Task | 6.26e-08 | 3.13e-08 | 3.32e-08 | 1.19e-08 | 0.46 | 74.74*** | .44 |
| Gambling Task | 10.22e-08 | 7.89e-08 | 5.35e-08 | 2.37e-08 | 3.15 | 38.07*** | .28 |
| Motor Task | 8.59e-08 | 6.68e-08 | 2.48e-08 | 1.19e-08 | 4.47* | 74.30*** | .43 |

*Note.* This table demonstrates means and standard deviations of measures for stability of the stable and instable state, as well as corresponding test statistics of repeated measures ANOVA significance testing. Means and standard deviations are rounded to increase comparability for the reader.

[a] CV = covariate, here mean difference of activation between the stable and instable state

*** $p < 0.001$, ** $p < 0.01$, * $p < 0.05$

**Table S16**

*Results of repeated measures ANOVA comparing control energy between exclusion of individual level rich club members and a size-matched set of semi-randomly selected regions*

| Measures | RC excluded | | Random excluded | | CV | Main effect | |
|---|---|---|---|---|---|---|---|
| | *M* | *SD* | *M* | *SD* | *F*(1,96) | *F*(1,96) | $\eta^2$ |
| Energy stable -> instable | | | | | | | |
| Emotion Task | 2.57e+07 | 1.58e+07 | 5.48e+07 | 3.00e+07 | 43.10*** | 17.24*** | .15 |
| Working Memory Task | 3.05e+07 | 1.85e+07 | 4.42e+07 | 1.73e+07 | 2.41 | 16.41*** | .15 |
| Language Task | 3.41e+07 | 1.85e+07 | 7.06e+07 | 2.57e+07 | 1.10 | 105.01*** | .52 |
| Relational Task | 3.52e+07 | 2.35e+07 | 5.19e+07 | 2.63e+07 | 0.00 | 20.43*** | .18 |
| Social Task | 2.91e+07 | 2.10e+07 | 5.05e+07 | 2.33e+07 | 8.05** | 41.72*** | .30 |
| Gambling Task | 1.80e+07 | 1.24e+07 | 2.73e+07 | 1.09e+07 | 0.95 | 27.56*** | .22 |
| Motor Task | 3.83e+07 | 3.66e+07 | 10.28e+07 | 5.63e+07 | 0.61 | 72.27*** | .43 |
| Energy instable -> stable | | | | | | | |
| Emotion Task | 2.03e+07 | 1.67e+07 | 3.03e+07 | 1.39e+07 | 1.77 | 10.72** | .10 |
| Working Memory Task | 2.22e+07 | 1.75e+07 | 3.17e+07 | 1.33e+07 | 0.94 | 13.05*** | .12 |
| Language Task | 3.66e+07 | 2.03e+07 | 6.72e+07 | 2.41e+07 | 0.01 | 123.43*** | .56 |
| Relational Task | 2.63e+07 | 1.57e+07 | 4.28e+07 | 2.83e+07 | 14.49*** | 3.75 | .04 |
| Social Task | 2.20e+07 | 1.30e+07 | 3.21e+07 | 1.77e+07 | 12.91*** | 6.20* | .06 |
| Gambling Task | 1.64e+07 | 1.16e+07 | 2.48e+07 | 1.04e+07 | 1.26 | 28.30*** | .23 |
| Motor Task | 0.82e+07 | 0.77e+07 | 2.02e+07 | 1.17e+07 | 20.29*** | 86.19*** | .47 |

*Note.* This table demonstrates means and standard deviations of measures for control energy for the transition between the stable and instable state, as well as corresponding test statistics of repeated measures ANOVA significance testing. Means and standard deviations are rounded to increase comparability for the reader.

[a] CV = covariate, here mean difference of activation between the stable and instable state

*** $p < 0.001$, ** $p < 0.01$, * $p < 0.05$

**Table S17**

*Results of repeated measures ANOVA comparing state stability between exclusion of individual level rich club members and a size-matched set of semi-randomly selected regions that have been matched regarding similarity of their connectivity profiles*

| Measures | RC excluded | | Random excluded | | CV | Main effect | |
|---|---|---|---|---|---|---|---|
| | *M* | *SD* | *M* | *SD* | *F*(1,96) | *F*(1,96) | $\eta^2$ |
| Stability stable state | | | | | | | |
| Emotion Task | 8.83e-08 | 5.89e-08 | 4.04e-08 | 1.50e-08 | 0.13 | 30.30*** | .24 |
| Working Memory Task | 8.26e-08 | 7.16e-08 | 3.70e-08 | 1.47e-08 | 0.08 | 23.10*** | .19 |
| Language Task | 5.59e-08 | 2.70e-08 | 2.54e-08 | 0.97e-08 | 5.96* | 118.61*** | .55 |
| Relational Task | 5.53e-08 | 3.80e-08 | 2.75e-08 | 0.99e-08 | 1.97 | 47.67*** | .33 |
| Social Task | 7.42e-08 | 4.21e-08 | 4.23e-08 | 1.76e-08 | 0.73 | 52.38*** | .35 |
| Gambling Task | 11.28e-08 | 9.95e-08 | 5.30e-08 | 2.32e-08 | 3.76 | 36.01*** | .27 |
| Motor Task | 22.42e-08 | 12.86e-08 | 8.81e-08 | 3.53e-08 | 0.39 | 84.41*** | .47 |
| Stability instable state | | | | | | | |
| Emotion Task | 6.78e-08 | 3.65e-08 | 2.98e-08 | 1.03e-08 | 0.28 | 50.99*** | .35 |
| Working Memory Task | 5.88e-08 | 3.87e-08 | 3.15e-08 | 1.16e-08 | 0.49 | 34.21*** | .26 |
| Language Task | 5.91e-08 | 3.29e-08 | 2.54e-08 | 1.11e-08 | 1.19 | 75.79*** | .44 |
| Relational Task | 4.71e-08 | 2.82e-08 | 2.49e-08 | 0.96e-08 | 1.78 | 51.02*** | .35 |
| Social Task | 6.26e-08 | 3.13e-08 | 3.32e-08 | 1.21e-08 | 0.63 | 79.67*** | .45 |
| Gambling Task | 10.22e-08 | 7.89e-08 | 4.96e-08 | 2.10e-08 | 3.22 | 42.34*** | .31 |
| Motor Task | 8.59e-08 | 6.68e-08 | 2.90e-08 | 1.59e-08 | 4.30* | 73.24*** | .43 |

*Note.* This table demonstrates means and standard deviations of measures for stability of the stable and instable state, as well as corresponding test statistics of repeated measures ANOVA significance testing. Means and standard deviations are rounded to increase comparability for the reader.

[a] CV = covariate, here mean difference of activation between the stable and instable state

*** $p < 0.001$, ** $p < 0.01$, * $p < 0.05$

**Table S18**

*Results of repeated measures ANOVA comparing control energy between exclusion of individual level rich club members and a size-matched set of semi-randomly selected regions that have been matched regarding similarity of their connectivity profiles*

| Measures | RC excluded | | Random excluded | | CV | Main effect | |
|---|---|---|---|---|---|---|---|
| | *M* | *SD* | *M* | *SD* | *F*(1,96) | *F*(1,96) | $\eta^2$ |
| Energy stable -> instable | | | | | | | |
| Emotion Task | 2.57e+07 | 1.58e+07 | 5.68e+07 | 3.10e+07 | 38.17*** | 23.51*** | .20 |
| Working Memory Task | 3.05e+07 | 1.85e+07 | 4.45e+07 | 1.70e+07 | 1.79 | 25.61*** | .21 |
| Language Task | 3.41e+07 | 1.85e+07 | 6.75e+07 | 2.54e+07 | 0.48 | 91.83*** | .49 |
| Relational Task | 3.52e+07 | 2.35e+07 | 5.46e+07 | 2.63e+07 | 0.00 | 31.82*** | .25 |
| Social Task | 2.91e+07 | 2.10e+07 | 4.89e+07 | 2.18e+07 | 4.14* | 72.34*** | .43 |
| Gambling Task | 1.80e+07 | 1.24e+07 | 2.87e+07 | 1.14e+07 | 3.42 | 40.78*** | .30 |
| Motor Task | 3.83e+07 | 3.66e+07 | 8.45e+07 | 4.78e+07 | 2.00 | 61.93*** | .39 |
| Energy instable -> stable | | | | | | | |
| Emotion Task | 2.03e+07 | 1.67e+07 | 3.39e+07 | 1.49e+07 | 0.89 | 26.19*** | .21 |
| Working Memory Task | 2.22e+07 | 1.75e+07 | 3.41e+07 | 1.47e+07 | 5.98* | 25.16*** | .21 |
| Language Task | 3.66e+07 | 2.03e+07 | 6.56e+07 | 2.55e+07 | 0.14 | 111.44*** | .54 |
| Relational Task | 2.63e+07 | 1.57e+07 | 4.53e+07 | 2.60e+07 | 12.98*** | 12.86*** | .12 |
| Social Task | 2.20e+07 | 1.30e+07 | 3.36e+07 | 1.84e+07 | 4.14* | 15.71*** | .14 |
| Gambling Task | 1.64e+07 | 1.16e+07 | 2.61e+07 | 1.13e+07 | 0.62 | 46.62*** | .33 |
| Motor Task | 0.82e+07 | 0.77e+07 | 1.82e+07 | 1.02e+07 | 8.84* | 86.21*** | .47 |

*Note.* This table demonstrates means and standard deviations of measures for control energy for the transition between the stable and instable state, as well as corresponding test statistics of repeated measures ANOVA significance testing. Means and standard deviations are rounded to increase comparability for the reader.

[a] CV = covariate, here mean difference of activation between the stable and instable state

*** $p < 0.001$, ** $p < 0.01$, * $p < 0.05$

**Table S19**

*Results of repeated measures ANOVA comparing state stability between exclusion of individual level rich club members and a size-matched set of semi-randomly selected regions that have been matched regarding total amount of connections*

| Measures | RC excluded | | Random excluded | | CV | Main effect | |
|---|---|---|---|---|---|---|---|
| | *M* | *SD* | *M* | *SD* | *F*(1,96) | *F*(1,96) | η[2] |
| Stability stable state | | | | | | | |
| Emotion Task | 8.83e-08 | 5.89e-08 | 5.27e-08 | 1.90e-08 | 0.33 | 16.13*** | .14 |
| Working Memory Task | 8.26e-08 | 7.16e-08 | 4.62e-08 | 1.81e-08 | 0.05 | 15.16*** | .14 |
| Language Task | 5.59e-08 | 2.70e-08 | 3.12e-08 | 1.16e-08 | 5.02* | 87.40*** | .48 |
| Relational Task | 5.53e-08 | 3.80e-08 | 3.38e-08 | 1.24e-08 | 1.89 | 33.33*** | .26 |
| Social Task | 7.42e-08 | 4.21e-08 | 5.07e-08 | 2.08e-08 | 0.58 | 32.54*** | .25 |
| Gambling Task | 11.28e-08 | 9.95e-08 | 6.69e-08 | 3.01e-08 | 3.26 | 24.36*** | .20 |
| Motor Task | 22.42e-08 | 12.86e-08 | 11.23e-08 | 4.63e-08 | 0.37 | 69.04*** | .42 |
| Stability instable state | | | | | | | |
| Emotion Task | 6.78e-08 | 3.65e-08 | 3.80e-08 | 1.31e-08 | 0.73 | 28.82*** | .23 |
| Working Memory Task | 5.88e-08 | 3.87e-08 | 3.87e-08 | 1.47e-08 | 0.24 | 20.25*** | .17 |
| Language Task | 5.91e-08 | 3.29e-08 | 3.08e-08 | 1.25e-08 | 1.23 | 61.52*** | .39 |
| Relational Task | 4.71e-08 | 2.82e-08 | 3.05e-08 | 1.20e-08 | 1.80 | 32.43*** | .25 |
| Social Task | 6.26e-08 | 3.13e-08 | 3.99e-08 | 1.43e-08 | 1.03 | 57.52*** | .37 |
| Gambling Task | 10.22e-08 | 7.89e-08 | 6.25e-08 | 2.60e-08 | 3.12 | 28.33*** | .23 |
| Motor Task | 8.59e-08 | 6.68e-08 | 3.72e-08 | 2.00e-08 | 3.97* | 60.34*** | .39 |

*Note.* This table demonstrates means and standard deviations of measures for stability of the stable and instable state, as well as corresponding test statistics of repeated measures ANOVA significance testing. Means and standard deviations are rounded to increase comparability for the reader.

[a] CV = covariate, here mean difference of activation between the stable and instable state

*** $p < 0.001$, ** $p < 0.01$, * $p < 0.05$

**Table S20**

*Results of repeated measures ANOVA comparing control energy between exclusion of individual level rich club members and a size-matched set of semi-randomly selected regions that have been matched regarding total amount of connections*

| Measures | RC excluded | | Random excluded | | CV | Main effect | |
|---|---|---|---|---|---|---|---|
| | *M* | *SD* | *M* | *SD* | *F*(1,96) | *F*(1,96) | $\eta^2$ |
| Energy stable -> instable | | | | | | | |
| Emotion Task | 2.57e+07 | 1.58e+07 | 4.42e+07 | 2.31e+07 | 36.33*** | 9.09* | .09 |
| Working Memory Task | 3.05e+07 | 1.85e+07 | 3.73e+07 | 1.49e+07 | 2.80 | 3.79 | .04 |
| Language Task | 3.41e+07 | 1.85e+07 | 5.61e+07 | 2.10e+07 | 0.54 | 62.44*** | .39 |
| Relational Task | 3.52e+07 | 2.35e+07 | 4.48e+07 | 2.01e+07 | 0.06 | 12.55*** | .12 |
| Social Task | 2.91e+07 | 2.10e+07 | 4.03e+07 | 1.72e+07 | 1.65 | 26.27*** | .21 |
| Gambling Task | 1.80e+07 | 1.24e+07 | 2.27e+07 | 0.91e+07 | 0.36 | 8.88** | .08 |
| Motor Task | 3.83e+07 | 3.66e+07 | 6.66e+07 | 3.69e+07 | 0.44 | 35.50*** | .27 |
| Energy instable -> stable | | | | | | | |
| Emotion Task | 2.03e+07 | 1.67e+07 | 2.56e+07 | 1.13e+07 | 2.05 | 2.74 | .03 |
| Working Memory Task | 2.22e+07 | 1.75e+07 | 2.73e+07 | 1.16e+07 | 2.03 | 3.25 | .03 |
| Language Task | 3.66e+07 | 2.03e+07 | 5.38e+07 | 2.04e+07 | 0.06 | 42.64*** | .31 |
| Relational Task | 2.63e+07 | 1.57e+07 | 3.66e+07 | 2.03e+07 | 12.00*** | 3.29 | .03 |
| Social Task | 2.20e+07 | 1.30e+07 | 2.76e+07 | 1.35e+07 | 2.76 | 4.92* | .05 |
| Gambling Task | 1.64e+07 | 1.16e+07 | 2.09e+07 | 0.93e+07 | 0.08 | 12.32*** | .11 |
| Motor Task | 0.82e+07 | 0.77e+07 | 1.45e+07 | 0.89e+07 | 9.17* | 45.67*** | .32 |

*Note.* This table demonstrates means and standard deviations of measures for control energy for the transition the stable and instable state, as well as corresponding test statistics of repeated measures ANOVA significance testing. Means and standard deviations are rounded to increase comparability for the reader.

[a] CV = covariate, here mean difference of activation between the stable and instable state

*** $p < 0.001$, ** $p < 0.01$, * $p < 0.05$

**Table S21**

*Results of repeated measures ANOVA comparing control energy between exclusion of rich club members and a size-matched set of semi-randomly selected regions when transitioning between states belonging to different tasks*

| From: | RC excluded | | Random excluded | | CV | Main effect | |
|---|---|---|---|---|---|---|---|
| | *M* | *SD* | *M* | *SD* | *F*(1,96) | *F*(1,96) | $\eta^2$ |
| Emotion Task, to: | | | | | | | |
| Working Memory Task | 3.91e+7 | 2.61e+7 | 6.68e+7 | 4.26e+7 | 0.01 | 8.42* | .08 |
| Language Task | 5.95e+7 | 3.73e+7 | 11.86e+7 | 6.15e+7 | 0.42 | 61.48*** | .39 |
| Relational Task | 4.59e+7 | 1.80e+7 | 6.22e+7 | 3.33e+7 | 0.30 | 1.88 | .02 |
| Social Task | 4.29e+7 | 2.59e+7 | 7.09e+7 | 4.20e+7 | 12.54*** | 3.54 | .04 |
| Gambling Task | 4.29e+7 | 2.55e+7 | 7.24e+7 | 4.55e+7 | 1.12 | 26.81*** | .22 |
| Motor Task | 6.03e+7 | 3.30e+7 | 10.22e+7 | 5.45e+7 | 10.45** | 72.59*** | .43 |
| Working Memory Task, to: | | | | | | | |
| Emotion Task | 5.37e+7 | 2.72e+7 | 6.29e+7 | 2.60e+7 | 24.44*** | 17.38*** | .15 |
| Language Task | 8.22e+7 | 3.95e+7 | 11.96e+7 | 4.07e+7 | 0.12 | 47.64*** | .33 |
| Relational Task | 4.04e+7 | 2.04e+7 | 4.84e+7 | 2.26e+7 | 18.90*** | 1.32 | .01 |
| Social Task | 5.24e+7 | 2.48e+7 | 6.50e+7 | 2.46e+7 | 11.64*** | 7.37** | .07 |
| Gambling Task | 4.85e+7 | 2.26e+7 | 5.78e+7 | 2.23e+7 | 4.75* | 28.86*** | .23 |
| Motor Task | 7.50e+7 | 3.27e+7 | 9.58e+7 | 3.38e+7 | 1.20 | 49.81*** | .34 |
| Language Task, to: | | | | | | | |
| Emotion Task | 7.55e+7 | 4.73e+7 | 14.16e+7 | 5.74e+7 | 28.61*** | 19.01*** | .17 |
| Working Memory Task | 8.29e+7 | 4.86e+7 | 14.80e+7 | 5.56e+7 | 1.31 | 15.13*** | .14 |
| Relational Task | 8.60e+7 | 4.89e+7 | 15.18e+7 | 5.31e+7 | 1.48 | 9.20** | .09 |
| Social Task | 7.59e+7 | 4.43e+7 | 14.15e+7 | 5.05e+7 | 13.84*** | 5.32* | .05 |
| Gambling Task | 8.16e+7 | 4.65e+7 | 14.15e+7 | 5.20e+7 | 3.03 | 21.52*** | .18 |
| Motor Task | 2.72e+7 | 1.59e+7 | 5.48e+7 | 2.24e+7 | 1.27 | 53.86*** | .36 |
| Relational Task, to: | | | | | | | |
| Emotion Task | 6.71e+7 | 3.04e+7 | 7.65e+7 | 3.32e+7 | 1.54 | 22.11*** | .19 |
| Working Memory Task | 5.71e+7 | 3.00e+7 | 6.80e+7 | 4.57e+7 | 0.29 | 6.96** | .07 |
| Language Task | 10.16e+7 | 4.49e+7 | 14.28e+7 | 5.56e+7 | 3.52 | 69.79*** | .42 |
| Social Task | 6.64e+7 | 2.98e+7 | 8.43e+7 | 3.77e+7 | 15.41*** | 12.20*** | .11 |
| Gambling Task | 6.51e+7 | 3.22e+7 | 7.73e+7 | 4.40e+7 | 0.65 | 42.18*** | .31 |
| Motor Task | 9.66e+7 | 4.36e+7 | 11.91e+7 | 5.17e+7 | 0.33 | 78.65*** | .45 |
| Social Task, to: | | | | | | | |
| Emotion Task | 5.00e+7 | 2.56e+7 | 6.76e+7 | 2.98e+7 | 2.43 | 25.81*** | .21 |
| Working Memory Task | 4.55e+7 | 2.62e+7 | 6.69e+7 | 3.05e+7 | 1.16 | 6.29* | .06 |
| Language Task | 6.70e+7 | 3.34e+7 | 11.53e+7 | 4.33e+7 | 0.06 | 42.72*** | .31 |
| Relational Task | 4.27e+7 | 2.23e+7 | 6.67e+7 | 2.71e+7 | 1.98 | 6.66* | .06 |
| Gambling Task | 5.23e+7 | 2.88e+7 | 7.34e+7 | 3.09e+7 | 0.96 | 22.30*** | .19 |
| Motor Task | 6.89e+7 | 3.59e+7 | 9.82e+7 | 3.97e+7 | 0.64 | 60.79*** | .39 |
| Gambling Task, to: | | | | | | | |
| Emotion Task | 2.83e+7 | 1.52e+7 | 3.87e+7 | 1.58e+7 | 34.17*** | 11.28** | .11 |
| Working Memory Task | 1.95e+7 | 1.38e+7 | 2.93e+7 | 1.30e+7 | 2.30 | 3.78 | .04 |
| Language Task | 5.26e+7 | 2.41e+7 | 8.46e+7 | 2.92e+7 | 0.91 | 51.05*** | .35 |
| Relational Task | 1.92e+7 | 1.11e+7 | 2.93e+7 | 1.32e+7 | 9.70*** | 0.01 | .00 |
| Social Task | 3.09e+7 | 1.63e+7 | 4.30e+7 | 1.58e+7 | 16.28*** | 2.28 | .02 |
| Motor Task | 4.24e+7 | 2.01e+7 | 5.73e+7 | 2.08e+7 | 0.15 | 62.35*** | .39 |

| | | | | | | | |
|---|---|---|---|---|---|---|---|
| Motor Task, to: | | | | | | | |
| Emotion Task | 2.52e+7 | 1.55e+7 | 4.90e+7 | 2.18e+7 | 29.27*** | 13.22*** | .12 |
| Working Memory Task | 2.62e+7 | 1.33e+7 | 4.79e+7 | 1.86e+7 | 1.79 | 7.29** | .07 |
| Language Task | 7.65e+7 | 4.67e+7 | 13.07e+7 | 4.94e+7 | 0.22 | 55.38*** | .37 |
| Relational Task | 3.11e+7 | 1.59e+7 | 5.17e+7 | 1.92e+7 | 1.70 | 4.29* | .04 |
| Social Task | 2.70e+7 | 1.50e+7 | 4.83e+7 | 1.93e+7 | 13.55*** | 3.46 | .03 |
| Gambling Task | 2.20e+7 | 1.14e+7 | 3.79e+7 | 1.58e+7 | 0.02 | 26.73*** | .22 |

*Note.* This table demonstrates means and standard deviations of measures for control energy for the transition between two states belonging to different tasks, as well as corresponding test statistics of repeated measures ANOVA significance testing. Means and standard deviations are rounded to increase comparability for the reader.

[a] CV = covariate, here mean difference of activation between state A and state B

*** $p < 0.001$, **$p < 0.01$, *$p < 0.05$

**Table S22**

*Mean rank of regional contribution per resting state network and over all stability and energy measures, respectively*

| | stability | | energy | |
|---|---|---|---|---|
| | $M_{rank(emp)}$ | $M_{rank(rand)}$ | $M_{rank(emp)}$ | $M_{rank(rand)}$ |
| Yeo 1 – central visual | 60.10*** | 109.67 | 67.65*** | 109.82 |
| Yeo 2 – peripheral visual | 103.47 | 110.23 | 106.34 | 110.22 |
| Yeo 3 – dorsal somatomotor | 132.43* | 109.89 | 131.77** | 109.85 |
| Yeo 4 – ventral somatomotor | 105.03 | 110.07 | 105.83 | 110.05 |
| Yeo 5 – posterior dorsal attention | 103.99 | 109.88 | 108.05 | 109.88 |
| Yeo 6 – somatomotor association | 118.44 | 110.06 | 119.02 | 110.02 |
| Yeo 7 – posterior ventral attention | 117.77 | 109.88 | 117.97 | 109.89 |
| Yeo 8 – anterior ventral attention | 114.62 | 109.76 | 111.56 | 109.86 |
| Yeo 9 – medial temporal-limbic | 119.81 | 110.15 | 116.96 | 110.15 |
| Yeo 10 – orbitofrontal-limbic | 98.18 | 110.08 | 93.58 | 110.05 |
| Yeo 11 – posterior frontoparietal | 137.47* | 109.88 | 135.76* | 109.82 |
| Yeo 12 – ventro-lateral frontoparietal | 91.73* | 110.24 | 93.67* | 110.17 |
| Yeo 13 – dorso-lateral frontoparietal | 124.21* | 109.69 | 119.73 | 109.82 |
| Yeo 14 – lateral temporal DMN | 113.65 | 109.80 | 114.21 | 109.91 |
| Yeo 15 – ventral DMN | 117.22 | 110.62 | 116.09 | 110.40 |
| Yeo 16 – dorsal DMN | 113.08 | 110.20 | 112.80 | 110.14 |
| Yeo 17 – lateral DMN | 117.55 | 110.27 | 114.37 | 110.21 |

*Note.* This table demonstrates the mean empirical rank per resting state network regarding regional contribution to stability and energy measures; empirical values were compared to a null distribution in which regional rank was permutated 10000-fold before calculating the mean per network. Significance values are based on this comparison, and the mean randomized rank per network is included here to indicate direction of effect. Means and standard deviations are rounded to increase comparability for the reader.

DMN = default mode network

*** $p < 0.001$, **$p < 0.01$, *$p < 0.05$
